# Individualized morphometric-similarity deviations in autism linked to cortical hierarchy and microarchitecture

**DOI:** 10.1101/2025.10.09.680886

**Authors:** Makliya Mamat, Yiyong Chen, Lin Li

## Abstract

Autism spectrum disorder (ASD) is marked by profound neurobiological heterogeneity, yet it remains unclear whether atypical brain organization reflects a diffuse low-amplitude pattern shared broadly across individuals or distinct spatially specific deviations that vary from person to person. Resolving this question requires methods that move beyond group averages to map individualized cortical atypicality against normative expectations. We built subject-level morphometric similarity networks in which edges quantify multivariate morphometric similarity between cortical regions. We then applied hierarchical Bayesian regression (HBR) normative models to estimate region-wise deviations from age- and sex-adjusted norms while accounting for site variation. Individuals with ASD carried a greater burden of extreme regional deviations, yet those deviations were focal and idiosyncratic rather than uniformly distributed. The spatial pattern of deviation aligned with canonical cortical hierarchies, shifting similarity toward sensory and visual poles and away from association cortex. Moreover, edge-level testing identified a sparse reconfiguration concentrated in occipito-temporal and cross-network connections. Clustering of individual deviation maps identified two robust subgroups that differ in the global sign of deviation and in their cognitive and molecular correlates. These results show that cortical atypicality in ASD is constrained by brain gradients, expressed in subgroup-specific forms, and linked to distinct biological and cognitive axes.

## 1 Introduction

Autism spectrum disorder (ASD) is characterized by wide variation in cognition, behavior, and neurobiology[1, 2, 3]. This pervasive heterogeneity frustrates attempts to find a single “ASD brain”[4, 5, 6] because conventional case-control studies often report inconsistent or small effects as group averages obscure focal, person-specific deviations [7, 8, 9]. Hence, a core question for neuroimaging studies of ASD is not only whether the brain differs on average, but how individual deviations are distributed across the cortical sheet — whether they reflect a low-amplitude pattern shared widely across affected people or instead emerge as large, spatially specific departures in a minority of individuals.

Normative modeling reframes this question. Rather than asking whether means differ, normative methods estimate an expected value for each brain measure conditional on age, sex and other covariates and then quantify how each person departs from that expectation[10, 11]. When applied to developmental and clinical cohorts, this shift in perspective often reveals that nominal group effects are driven by a minority of individuals with extreme, spatially focal deviations while many affected people fall within the expected range [12, 13, 14, 11]. Applications in ASD also illustrate this point. For example, Gaussian process based norms showed highly idiosyncratic cortical thickness deviations in ASD despite weak case–control differences [13]. Subsequent work using large, multisite samples confirmed that normative outliers account for much of the tiny average effects seen in standard analyses [15]. Hierarchical Bayesian regression (HBR) implementations extend these ideas to multisite datasets by modelling site and scanner variation as random effects and by borrowing statistical strength across cohorts, producing calibrated per-region z scores that explicitly reflect uncertainty[16, 17, 18, 19]. In practice these calibrated deviation maps let us ask different questions than conventional group tests. They identify where and in whom the cortex diverges from expected developmental trajectories and therefore serve as a principled substrate for stratification and for testing whether deviations map onto canonical cortical hierarchies.

A meaningful interpretation of these deviations requires a spatial framework. The cortex is organized along reproducible, multiscale axes that reflect laminar structure, microstructure, gene expression and connectivity [20]. Macroscopically, low-dimensional connectivity gradients capture a sensory–transmodal hierarchy, with primary somatosensory/motor and visual areas at one extreme and the default-mode network (DMN) at the other[21, 22]. In ASD, this sensory–transmodal hierarchy appears perturbed [23, 24]. For instance, Hong et al. found reduced differentiation between sensory and higher-order systems, characterizing it as atypical connectivity transitions between unimodal sensory areas and transmodal DMN regions.[25]. Lee et al. also found reduced differentiation between sensory and transmodal systems and atypical maturation of principal gradients has been reported in ASD individuals [26]. They observed delayed development of this hierarchy in childhood and persistent impairments in DMN integration, implying that ASD brains follow a non-linear growth trajectory of cortical hierarchy. These findings suggest that ASD-related perturbations may be channeled along specific hierarchical axes rather than distributed uniformly across the cortex.

A related organizational scaffold is cortical chemoarchitecture, the spatial arrangement of neurotransmitter receptors and transporters across the cortical sheet [27]. Studies have shown that receptor densities vary systematically across the cortex, aligning with both structural and functional hierarchies [28, 29], and cortical areas share “receptor fingerprints” that delineate sensory versus association regions[30]. In practice, this means that an atypicality in ASD could associate with specific neurochemical systems (e.g. an excess of deviation in sensory cortex might implicate cholinergic or glutamatergic circuitry), whereas deviation in association cortex might implicate modulatory systems.

Structural similarity networks provide a principled, subject-level way to capture previously described multiscale cortical organization in a meaningful way [31]. In these networks each cortical region (node) is represented by a multifeature fingerprint (for example thickness, area, curvature) and cortico-cortical relationship (edge) reflect the similarity between region’ fingerprints, so that the connectome encodes multivariate affinities rather than independent univariate values[32, 33]. Anatomical homophily, the empirical regularity that areas similar in cytoarchitecture or molecular identity tend also to be similar in connectivity and function, gives this representation biological meaning and predicts that similarity matrices should align with histology and gene expression maps [34, 35, 36]. Indeed, early work showed that morphometric similarity networks recapitulate microscale cortical organization and transcriptomic gradients [32], and more recent methods that compare full multivariate distributions, such as Morphometric INverse Divergence (MIND)[33], improve reliability and increase correspondence to tract-tracing and transcriptomic benchmarks[33]. Crucially, structural similarity is defined at the single-subject level, which permits normative modeling of per-region atypicality and direct tests of whether individual deviation maps cluster or map onto canonical hierarchies. A recent study applying morphometric similarity in ASD has reported altered network topology, reduced global efficiency and atypical regional similarity in association cortices[37]. These suggest that individualized similarity deviations could capture the higher-order reconfiguration present in some autistic individuals.

In this study, we combined subject-wise MIND networks with HBR normative modeling in a large multisite cohort of ASD and typically developing (TD) individuals. We derived per-region, per-subject z-maps of cortical deviation, then examined whether these deviations are: (i) prevalent and focal at the individual level, (ii) nonrandomly distributed along canonical cortical hierarchies, and (iii) organized into reproducible subgroups with distinct molecular and cognitive profiles. This approach provides a principled framework to link individualized neuroanatomical deviations in ASD to cortical hierarchy and microarchitecture.

## 2 Results

### 2.1 Normative models reveal increased prevalence of extreme per-region deviation in ASD

To predict mean MIND values (nodal degree) across 308 cortical regions, we built HBR normative models that included B-spline basis functions for age, a fixed effect for sex, and random intercepts and slopes by site. The models were trained on an 80% site-stratified subset of TD participants and tested on the held-out TD and all ASD subjects. They showed stable performance, with a median explained variance of 0.15 (IQR 0.11–0.19) and well-calibrated test-set z-scores (Fig. S3, Fig. S4, Fig. S5).

Aggregating subject × region of interest (ROI) z-scores revealed significant differences in the prevalence of extreme deviation (|z| >2) between ASD and TD groups. ASD individuals exhibited higher rates of extreme deviations than TD controls in both positive (ASD: 3.80 ± 1.14%; TD: 3.05 ± 1.47%; Mann–Whitney *U* = 61,952, *p* = 1.42 × 10^−11^) and negative (ASD: 3.41 ± 1.15%; TD: 3.01 ± 1.51%; Mann–Whitney *U* = 55,242, *p* = 2.91 × 10^−5^) directions (Fig. 1A-C). Importantly, fewer than 12% of ASD individuals exhibited extreme deviations in any given brain region, highlighting that these MIND atypicalities are highly individualized and not a widespread characteristic across the ASD population within any single region.

**Figure 1.**
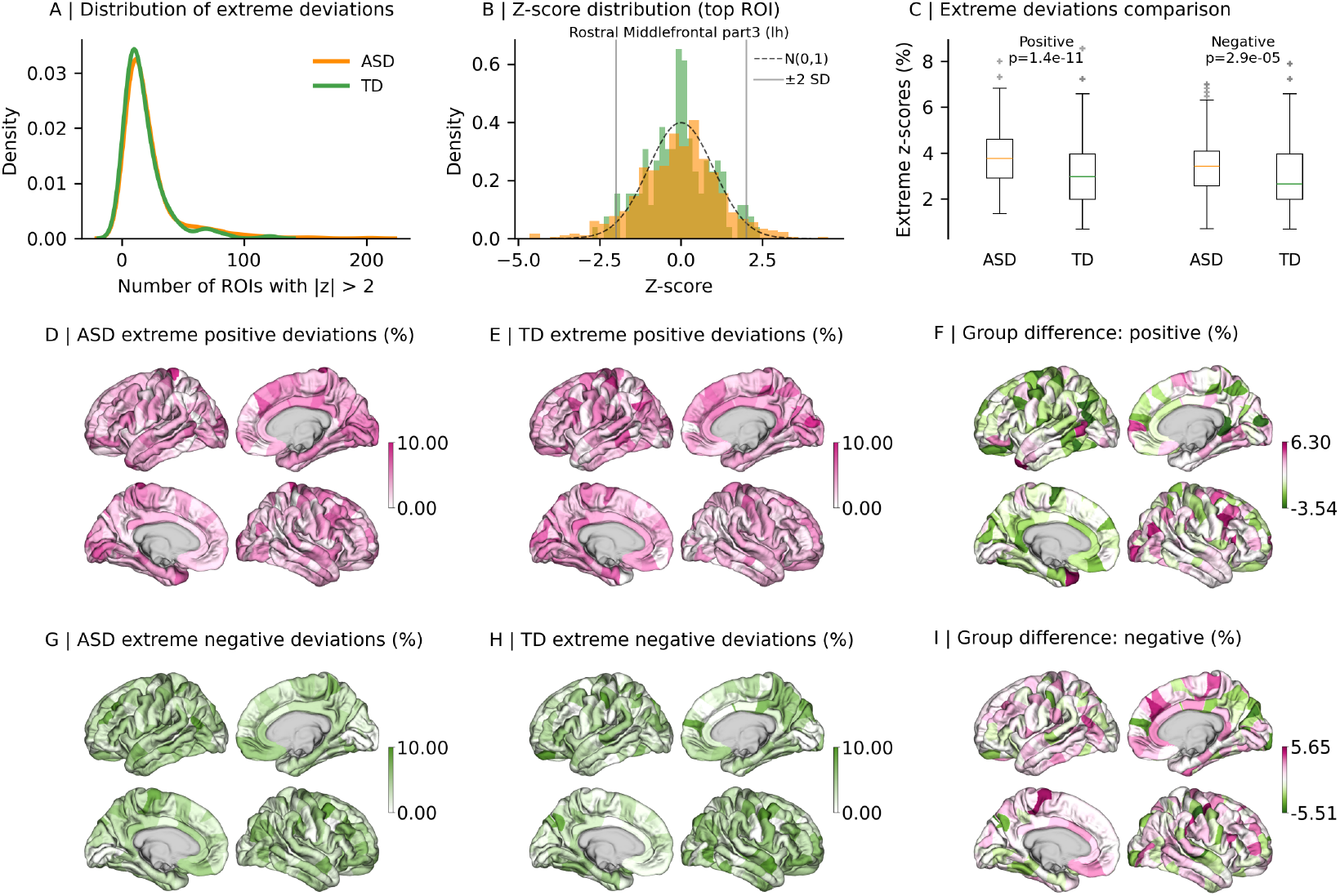
Anatomical distribution and prevalence of extreme normative deviations. Group-level characterization of suprathreshold deviations (*z >* 2). (A) Distribution of extreme deviations showing per-subject counts of extreme regional deviations in ASD (orange) and TD (green). (B) Z-score distribution for the ROI with highest extreme deviation prevalence, displaying individual z-scores for TD (green) and ASD (orange) groups with normal distribution reference (dashed line) and ±2 SD thresholds (gray lines). (C) Comparison of ROI-wise extreme percentages with nonparametric group tests (Mann–Whitney U). Region-wise prevalence maps for positive (D–F) and negative (G–I) extremes in ASD and TD, and the ASD–TD difference (percentage points) displayed on the cortical surface (diverging palette highlights excess prevalence in ASD).

Next, we explored the spatial distribution of atypical mean MIND patterns. Regions with the highest positive deviation differences (ASD–TD) included the left temporal pole (6.30%), left middle temporal cortex (5.30%), right caudal middle frontal cortex (4.65%), left pars orbitalis (4.48%), and right inferior parietal cortex (4.14%) (Fig. 1D-F). These findings suggest that temporal and frontal association regions show elevated morphometric similarity in ASD compared to normative references. In contrast, regions with the largest negative deviation differences included the left paracentral cortex (5.65%), left inferior parietal cortex (4.18%), right precentral cortex (3.77%), and left rostral middle frontal cortex (3.70%) (Fig. 1G-I).

Across the cohort, we found strong left–right correspondence and negligible case–control differences in bilateral homology. Inter-hemispheric correlations were robust in both groups (ASD: *r* = 0.311, *p <* 0.001; TD: *r* = 0.272, *p <* 0.001), indicating preserved inter-hemispheric structural symmetry in ASD. Inter-hemispheric asymmetry was minimal between groups (Cohen’s *d* = 0.035, *p <* 0.001), further supporting the preservation of bilateral structural organization in ASD (Fig. S8). These findings demonstrate that while ASD is associated with spatially focal deviations in MIND patterns, the fundamental inter-hemispheric structural organization remains intact.

In addition to these variability in extreme deviation prevalence, our findings also suggest that traditional case– control analyses may fail to capture the nuanced deviations observed in ASD (Fig. S9). Linear regression comparisons between ASD and TD groups, adjusting for age, sex, and site, revealed subtle group differences in only 2 of 308 regions, with effect sizes (Cohen’s *d* = 0.199 and *d* = 0.205) too small to be meaningful. These results reinforce our hypothesis that group-level comparisons are insufficient to detect the true spectrum of cortical reconfigurations in ASD. Indeed, our normative z-scores identified a subset of individuals with highly atypical brain patterns (Fig. S9A). Removing these outliers from the analysis eliminated all significant case–control differences, further suggesting that observed group differences are largely driven by a small subset of individuals with extreme atypicalities.

### 2.2 Morphometric deviations in ASD align with canonical zonal systems and functional gradients

We tested whether the spatial distribution of ASD-related morphometric deviations (operationalized as region-wise mean normative z-scores in the ASD group) is structured by canonical cortical systems or instead forms a diffuse perturbation. This question addresses two related hypotheses: (i) that deviations preferentially affect specific cytoarchitectonic/zonal classes (consistent with homophily principle), and (ii) that deviations are embedded within continuous macroscale gradients of cortical organization. To evaluate these hypotheses, we computed the ASD group mean z-score at each ROI and tested its relationship to canonical partitions: [38], von Economo cytoarchitectonic classes [39], the Yeo7 intrinsic networks [40], and canonical functional gradients [21].

Zone-level tests of the mean ASD z-score revealed systematic, regionally specific effects. Idiotypic and sensory zones showed significantly higher mean ASD z-scores (i.e., more positive deviations), whereas association and motor zones showed lower mean ASD z-scores (i.e., more negative deviations). Idiotypic cortex in the Mesulam taxonomy had higher mean z-scores (*t* = 2.739, *p*_FDR_ = 3.20 × 10^−2^), while von Economo association region 2 showed lower mean z-scores (*t* = −4.034, *p*_FDR_ = 2.31 × 10^−3^). Secondary sensory areas exhibited higher mean z-scores (*t* = 2.941, *p*_FDR_ = 2.14 × 10^−2^) and motor cortex showed lower mean z-scores (*t* = −3.014, *p*_FDR_ = 3.20 × 10^−2^). In the Yeo7 partition, the visual network had higher mean ASD z-scores (*t* = 3.566, *p*_FDR_ = 5.99 × 10^−3^) (Fig. 2A).

**Figure 2.**
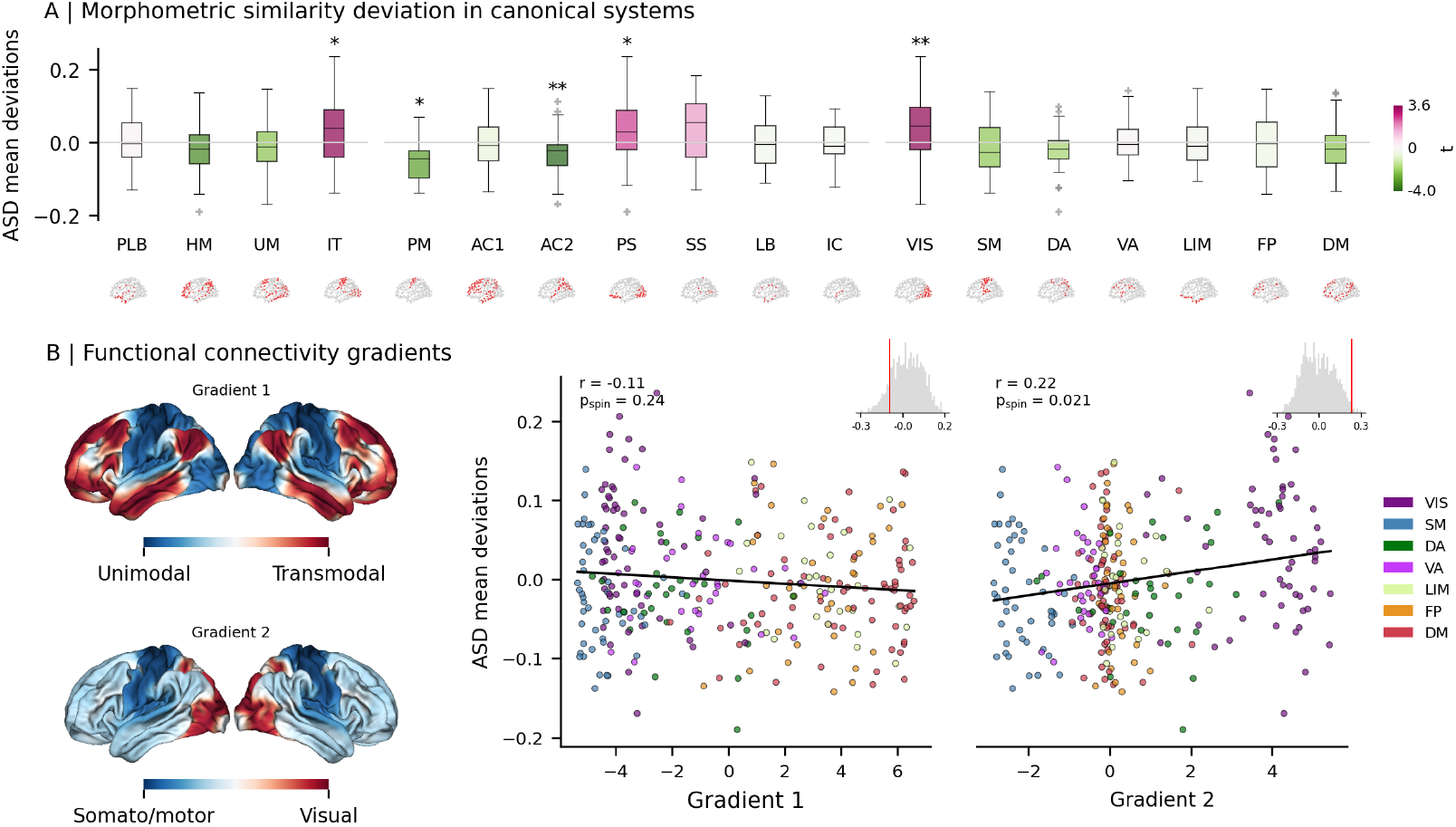
Alignment of ASD morphometric deviations with canonical zonal systems and functional gradients. (A) Distribution of ASD mean z-scores stratified by three cortical taxonomies: Mesulam laminar classes (left), von Economo cytoarchitectonic classes (middle), and Yeo7 intrinsic networks (right). Box summaries indicate central tendency and dispersion. Statistical significance is indicated by asterisks (** *p*_FDR_ *<* 0.01, * *p*_FDR_ *<* 0.05). (B) Left: surface renderings of canonical functional gradients. Right: scatterplots relating regional gradient values to ASD mean z-scores, colored by Yeo7 network. Histograms show spin-permutation null distributions (*n* = 10,000); empirical correlations and spin *p*-values are reported. Abbreviations: PLB = paralimbic, HM = heteromodal, UM = unimodal, IT = idiotypic, PM = primary motor, AC = association, PS = primary sensory, SS = secondary sensory, LB = limbic, IC = insula, VIS = visual, SM = somatomotor, DA = dorsal attention, VA = ventral attention, LIM = limbic, FP = frontoparietal, DM = default mode.

Pairwise zone contrasts confirmed a systematic redistribution of morphometric similarity between sensory and association territories. Of 48 zone–zone comparisons, 14 survived FDR correction (*p*_FDR_ *<* 0.05).Specifically, the visual and primary/secondary sensory regions exhibited stronger mean ASD z-scores relative to many association and heteromodal regions, and association/heteromodal regions (notably von Economo association 3) and motor cortex showed reduced mean z-scores relative to primary and secondary sensory regions (Fig. S10).

We then projected regional ASD mean z-scores onto canonical functional gradients. There was no significant association with the principal unimodal–to–transmodal axis (Gradient 1; *r* = − 0.11, p_spin_ = 0.24), but ASD mean z-scores were positively associated with the second canonical gradient (Gradient 2; *r* = 0.22, p_spin_ = 0.021) (Fig. 2B). Interpreting Gradient 2 spatially, regions nearer the visual pole (larger Gradient 2 values) tend to show more positive ASD mean z-scores, whereas regions nearer the somatomotor pole (smaller Gradient 2 values) tend to show smaller or negative deviations. This axis-specific reorganization implies a shift in the balance between local (sensory/visual) and distributed (association) similarity in ASD.

We also assessed whether the Gradient 2 alignment holds at the level of individual subjects. For each participant we correlated their regional z-map with Gradient 2; the mean subject-wise correlation was small but reliably positive (mean *r* = 0.014, 95% CI [0.005, 0.024]; one-sample *t*(586) = 3.47, *p* = 2.7 × 10^−3^; Cohen’s *d* = 0.12), indicating a subtle, group-level bias along Gradient 2 that is heterogeneously expressed across individuals (Fig. S11).

Together, these results indicate that ASD-related cortical deviations are not randomly distributed. Instead, deviations preferentially redistribute morphometric similarity from association/heteromodal cortex toward sensory and visual territories, and this reorganization is aligned with canonical cortical gradients rather than reflecting a uniform global shift.

### 2.3 Network-level reconfiguration of morphometric similarity in ASD

The regional deviations described above invite the question of whether atypicality in ASD also reshapes the broader organization of morphometric similarity networks. Unlike traditional structural connectomes, where edges denote anatomical tracts, MIND networks quantify similarity in morphometric profiles between cortical regions. In this framework, local deviations in morphometric profiles could propagate to alter distributed similarity coupling, producing a structured reconfiguration of the cortical network. To test this possibility, we applied the Network-Based Statistic (NBS) (52) to residualized MIND matrices after regressing out age, sex, and site effects. This analysis identified a significant subnetwork (5,000 permutations, *p*_FWE_ *<* 0.05) comprising 328 edges and 177 cortical nodes.

This subnetwork was not balanced in its composition. Most altered connections (259 edges) showed greater similarity in ASD than in TD, whereas a smaller fraction (69 edges) showed the reverse pattern (Fig. 4A–B). This asymmetry points to a relative strengthening of morphometric similarity among occipital and adjacent association regions, accompanied by more focal reductions in certain homotopic and association pairings. The most central nodes in this subnetwork, defined by their degree within the component, were located in the right and left lateral occipital cortex and the right entorhinal cortex, underscoring the prominence of occipito-temporal territories in the reorganization.

Visualization of the full edgewise contrast confirmed that the NBS result occupies a spatially structured subset of the matrix rather than reflecting a diffuse global shift. Clustered pockets of increased and decreased similarity were evident in the unthresholded maps, but only a small fraction of edges survived component-level correction, consistent with a localized pattern of network change (Fig. S12).

To better understand the functional implications of these network changes, we stratified the significant edges by Yeo7 functional networks. The visual system accounted for the largest number of altered couplings (214 edges), followed by default (112 edges), somatomotor (82 edges), ventral attention (61 edges), limbic (59 edges), and frontoparietal (52 edges), whereas the dorsal-attention system contributed the fewest (43 edges). Nearly 90% of significant edges (295 of 328) linked nodes across different networks, suggesting that the reconfiguration is driven primarily by altered between-network similarity rather than changes confined within a single system (Fig. 3C).

**Figure 3.**
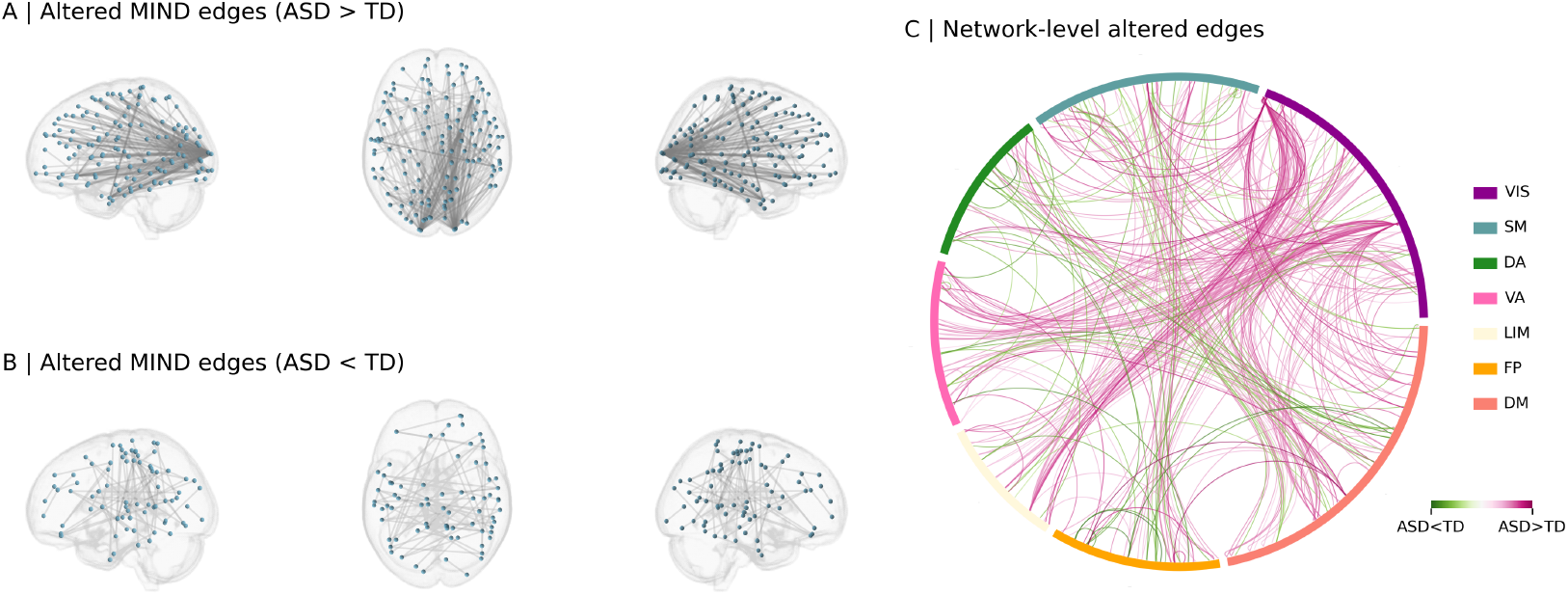
Network-level reconfiguration of morphometric similarity in ASD identified by NBS. Spatial rendering of edges within the NBS-identified component that show significantly greater similarity in ASD than TD (A) and significantly reduced similarity in ASD (B) (P_FDR_ < 0.05). (C) Chord diagram summarizing significant cross-network and within-network edges by Yeo7 partition; chord width encodes the number of altered edges and color indicates direction of change. The significant subnetwork is dominated by occipito-temporal/visual coupling and by cross-network reconfiguration.

**Figure 4.**
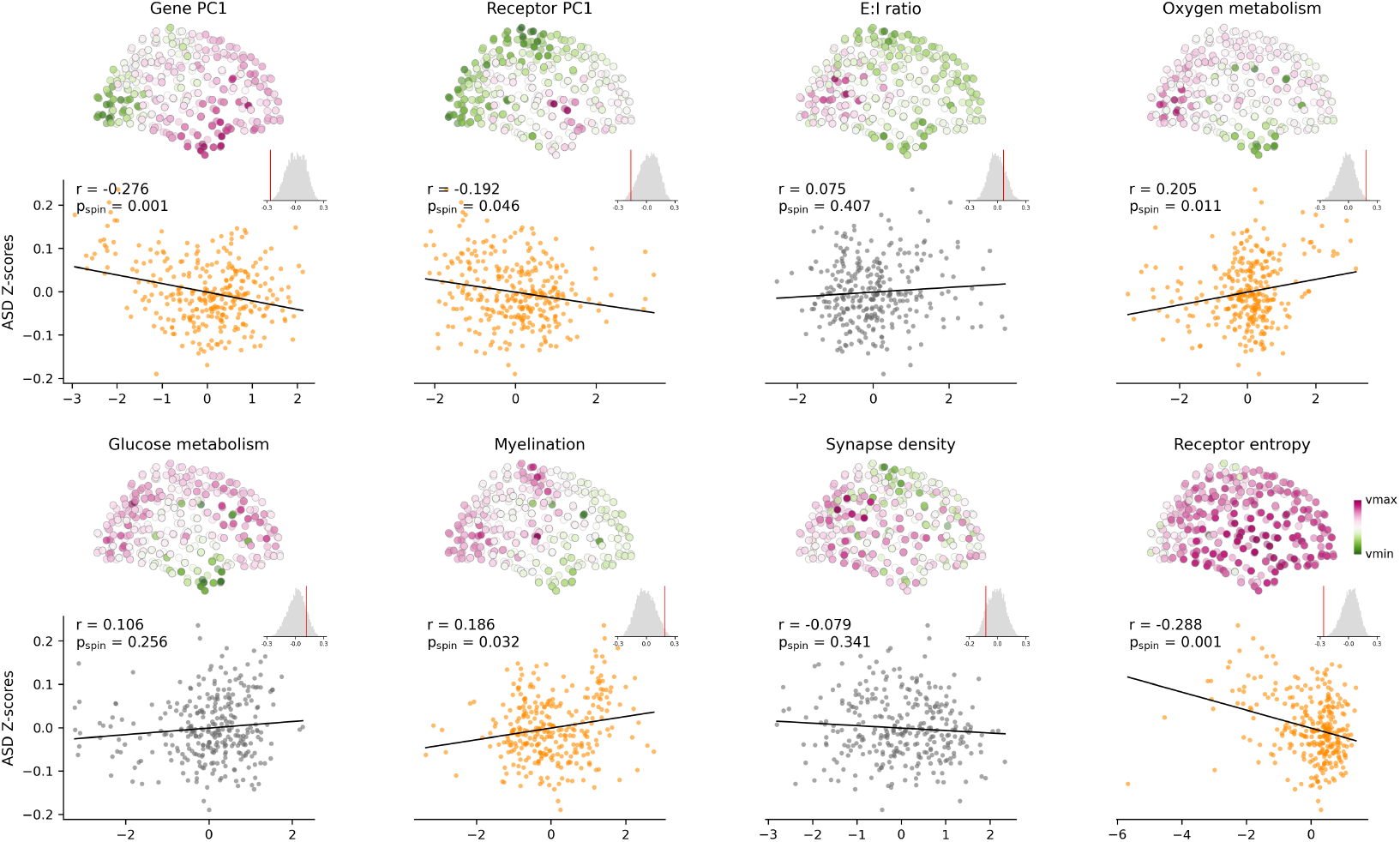
Coupling of ASD morphometric deviations to molecular and metabolic cortical maps. For each attribute (gene-expression PC1, receptor PC1, receptor entropy, E:I ratio, oxygen/glucose metabolism, myelination, synapse density), cortical maps are paired with scatterplots showing regional ASD mean z-scores versus the attribute. Points in orange indicate region-wise correlations that survive spin-permutation testing (p_spin_ < 0.05; 10,000 spins); gray points do not. Reported *r* and p_spin_ values quantify spatial associations.

### 2.4 Macroscale morphometric similarity deviations align with microscale cortical properties

We next asked whether macroscale morphometric deviations in ASD reflect intrinsic micro-architectural properties of cortical regions. Classic network models often assume regional homogeneity, but cortical areas differ markedly in their molecular and developmental programs. The homophily principle[31] suggests that macroscale connectivity is constrained by these microscale attributes. We therefore hypothesized that ASD-related deviations would follow systematic molecular and metabolic gradients rather than being randomly distributed across cortex.

To test this, we compared the spatial distribution of ASD mean z-scores with biologically informed cortical maps, including gene-expression PC1 (gene PC1), receptor PC1, excitatory–inhibitory (E:I) receptor balance, oxygen and glucose metabolism, myelination, synapse density, and receptor entropy (Fig. S13).

The results revealed selective associations with molecular and metabolic gradients (Fig. 4). Gene PC1 was negatively correlated with ASD deviations (*r* = − 0.276, pspin=0.001), indicating that regions with strong loadings on the dominant transcriptional axis show attenuated or negative morphometric changes. Similar negative correlations were observed for receptor PC1 (*r* = − 0.192, pspin=0.046) and receptor entropy (*r* = − 0.288, pspin=0.001), suggesting that areas with diverse receptor repertoires preferentially show reduced similarity in ASD. In contrast, oxygen metabolism (*r* = 0.205, pspin=0.011) and myelination (*r* = 0.186, p_spin_=0.032) were positively associated, pointing to relative increases in similarity within metabolically active and more myelinated territories.

Together, these findings suggest a structured anatomical ordering. That is, decreases in morphometric similarity cluster within transcriptomically defined association cortex, whereas increases concentrate in metabolically active sensory regions. This may suggests that the cortex’s intrinsic molecular and metabolic architecture constrains where morphometric reconfiguration in ASD is most likely to occur.

### 2.5 Divergent cortical–molecular fingerprints define ASD subgroups

Another important question motivating this study is whether individualized morphometric-similarity deviations in ASD reflect multiple, biologically meaningful phenotypes or a single weakly expressed pattern spread across many individuals. To address this, we clustered subject-level MIND deviation maps using a discriminative, case–control– aware algorithm and identified two stable subgroups (adjusted Rand index = 0.783) (Fig. 5A–D). The clusters are distinguished most simply by the global sign of deviation: ASD1 carries net positive regional z-scores (mean z = 0.412 ± 0.099), whereas ASD2 carries net negative z-scores (mean z = − 0.370 ± 0.114). This polarity provided a natural axis for asking whether subgroup differences map onto cortical hierarchy, molecular gradients, and functional systems.

**Figure 5.**
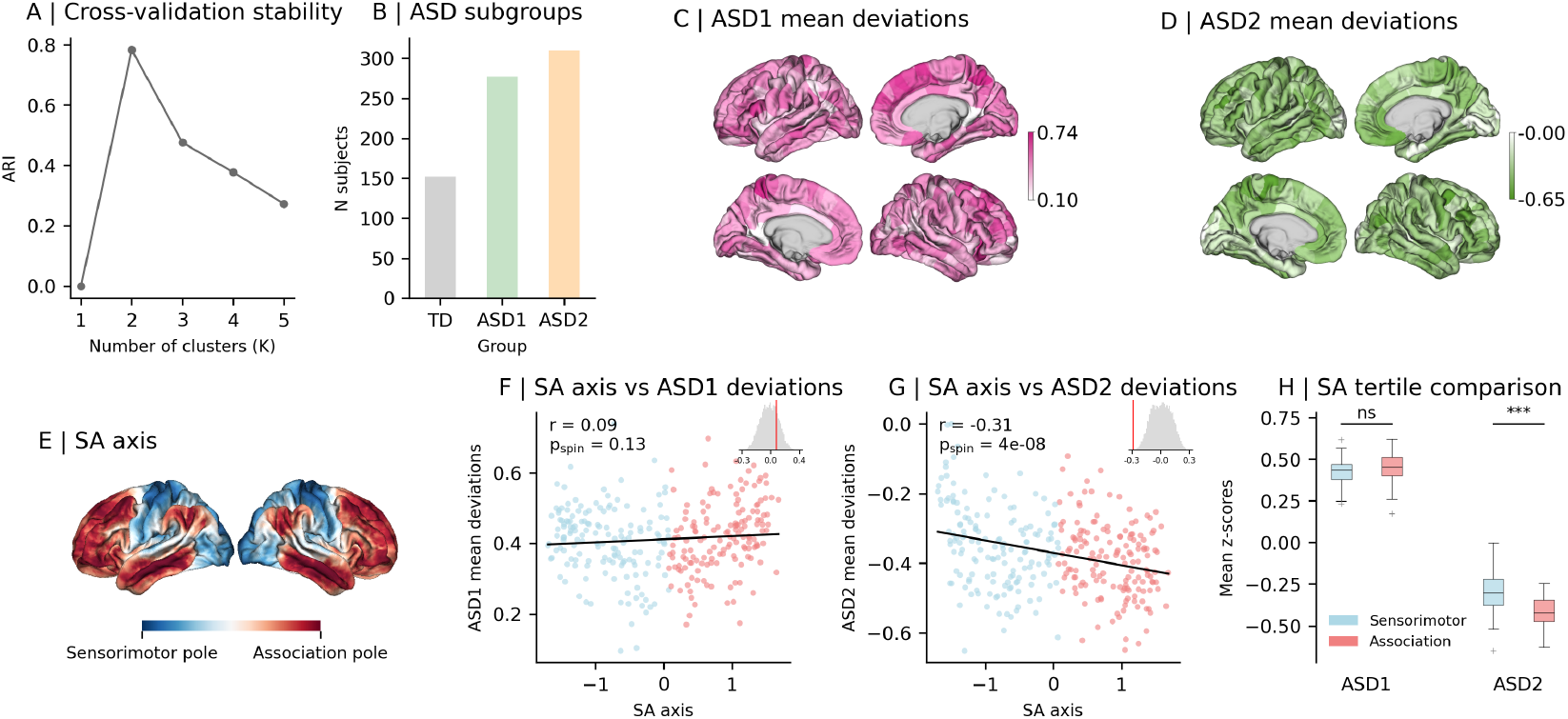
Two reproducible ASD subgroups with distinct hierarchical embeddings. (A–B) Cluster stability (adjusted Rand index) across cross-validation folds identifies an optimal *K* = 2 solution (maximum ARI = 0.783). (C–D) Surface maps of subgroup mean z-scores: ASD1 (net positive deviations) and ASD2 (net negative deviations). (E) Sensorimotor–association (S–A) axis displayed on the cortical surface (sensorimotor pole in blue; association pole in red). (F–G) Scatterplots and spin-permutation nulls (*n* = 10,000) relating S–A coordinate to subgroup mean deviations (ASD1: *r* = 0.085, p_spin_ = 0.14; ASD2: *r* = −0.306, p_spin_ = 4.1 × 10^−8^). (H) Tertile contrast (bottom 60 sensorimotor ROIs vs. top 60 association ROIs) quantifies subgroup divergence (ASD1: *t* = −1.96, p_spin_ = 0.053; ASD2: *t* = 5.49, p_spin_ = 2.38 × 10^−7^).

A central axis of cortical organization is the sensorimotor–association (SA) axis[41], which reflects a gradient from primary sensory and motor areas to complex association cortex(Fig. 5E). Projecting subgroup mean deviations onto the SA axis revealed a clear dissociation. ASD1 did not show a reliable bias along S–A (*r* = 0.085, p_spin_ = 0.14). By contrast, ASD2 aligned strongly toward the sensorimotor/visual pole (*r* = − 0.306, p_spin_ = 4.1 ×10^−8^) (Fig. 5F–G). A tertile contrast (bottom 60 sensorimotor parcels vs. top 60 association parcels) replicated this result (ASD1: *t* = − 1.96, p_spin_ = 0.053; ASD2: *t* = 5.49, p_spin_ = 2.38 × 10^−7^) (Fig. 6H). In short, ASD2 expresses a systematic redistribution of morphometric similarity away from association cortex and toward sensory territories; ASD1 lacks this consistent hierarchical bias.

**Figure 6.**
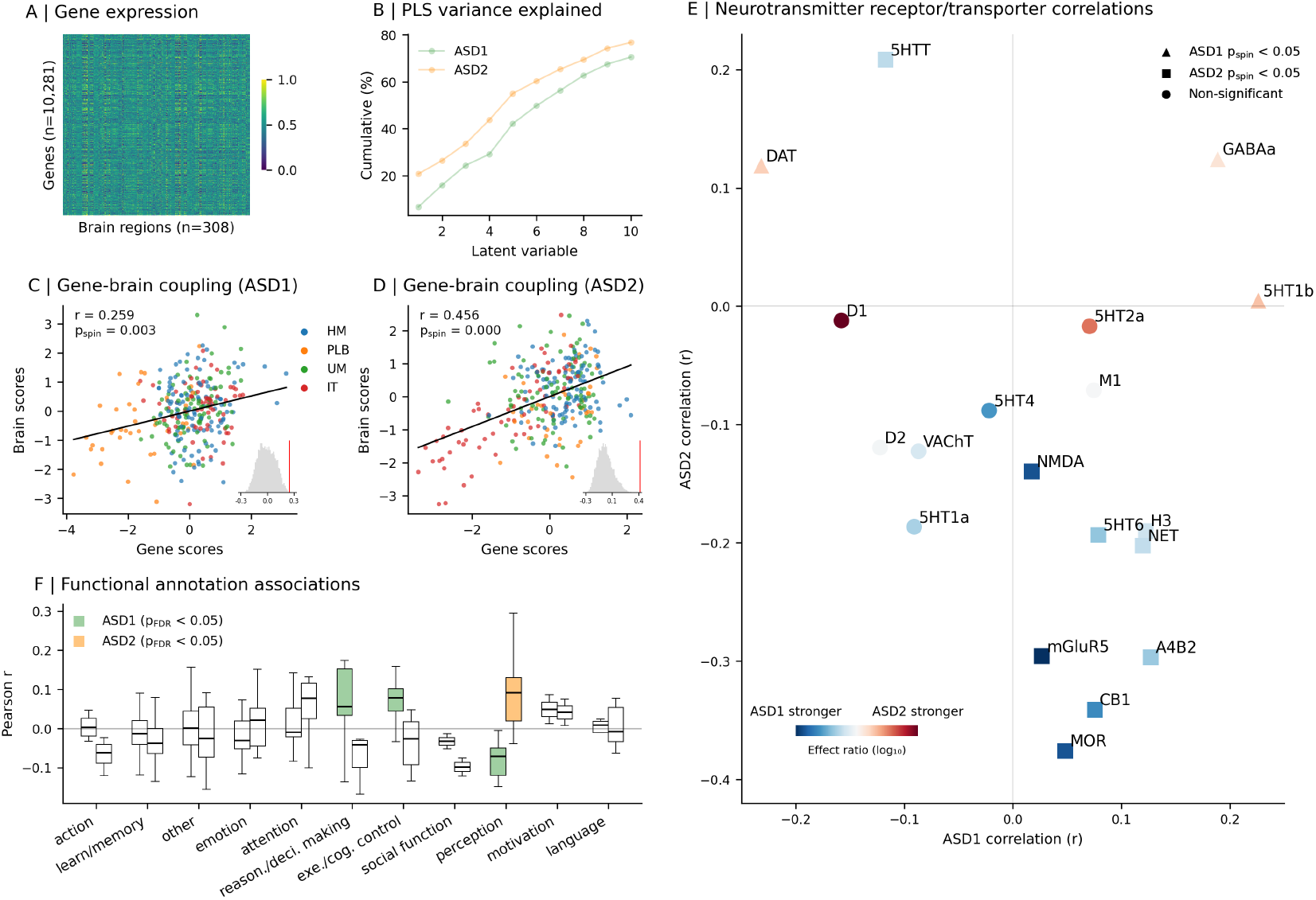
Multiscale molecular and functional correlates of subgroup-specific deviations. (A) Regional gene-expression heatmap (Allen Human Brain Atlas, genes × 308 regions) used for PLS regression. (B) Cumulative variance explained by PLS latent variables for ASD1 (green) and ASD2 (orange). (C–D) Gene–brain scatterplots for LV1: gene scores (PLS gene-side LV1) versus brain scores (PLS brain-side LV1), colored by Mesulam laminar class; insets show spin null distributions (*n* = 10,000) and empirical correlations. LV1 explains substantially more variance for ASD2 than ASD1. (E) Neurotransmitter receptor/transporter association plot. Each point represents a molecular target; marker shape indicates statistical significance of receptor-brain coupling (triangle = ASD1 p_spin_ < 0.05; square = ASD2 p_spin_ < 0.05; circle = non-significant). Marker color encodes the log_10_ effect-ratio of receptor-brain coupling strength: red indicates ASD2 shows stronger molecular coupling than ASD1, while blue indicates ASD1 shows stronger coupling than ASD2. The color intensity reflects the magnitude of the difference in coupling strength between subgroups. (F) Correlations with Neurosynth meta-analytic maps. Boxplots group correlations by functional domain (123 terms). Statistical testing used FDR correction across terms and categories.

To identify molecular substrates of the subgroup-specific deviation patterns we related regional gene expression to subtype deviation maps using partial least squares (PLS) regression and focused on the first latent variable (LV1)(Fig. 6A– B; Fig. S14A). LV1 showed robust brain–expression coupling for both subtypes, with spatial correlations *r* = 0.259 for ASD1 (p_spin_ = 0.003) and *r* = 0.456 for ASD2 (p_spin_ < 0.001). Bootstrap resampling confirmed stable gene loadings for LV1. Spatially, ASD1 LV1 weights concentrated on frontal and motor regions (positive) and insular/temporal areas (negative), whereas ASD2 LV1 weights placed positive emphasis on frontal/parietal parcels and negative emphasis on occipital/visual cortices (Fig. 6C–D). In distance-based train–test splits LV1 retained modest out-of-sample correspondence for ASD1 (test *r* ≈ 0.135 ± 0.092) and substantially higher correspondence for ASD2 (test *r* ≈ 0.420 ± 0.090), consistent with a tighter transcriptional constraint on the ASD2 phenotype than on ASD1 (Fig. S14B–D).

We then evaluated predictive models built from PET-derived receptor densities and from aggregated biological attributes to identify molecular drivers of the subgroup deviation patterns (Fig. S15, Fig. S16). Receptor-based models consistently outperformed attribute-based models (higher *R*^2^), with fits highest for ASD2 (receptor *R*^2^ ≈ 0.276; biological *R*^2^ ≈ 0.213), intermediate for pooled ASD, and lowest for ASD1 (receptor *R*^2^ ≈ 0.118; biological *R*^2^ ≈ 0.075)(Fig. **??**, Fig. S17). Dominance decomposition highlighted glucose metabolism and gene-PC1 as principal contributors for ASD2, whereas dopaminergic markers (DAT, D1) carried more weight for ASD1. Serotonergic markers (notably 5-HTT) also ranked highly for ASD2. Distance-based cross-validation confirmed higher spatial generalizability of receptor models for ASD2 (Fig. S17B–C). To pinpoint receptor-level drivers, we correlated each subtype’s regional deviation map with PET-derived receptor/transporter density maps and assessed significance with spin-permutation testing. ASD1 exhibited only a few significant associations (positive with 5-HT1b and GABAa; negative with DAT), while ASD2 showed a broader, statistically robust fingerprint (negative correlations with MOR, CB1, *α*4*β*2, and mGluR5, and a positive correlation with 5-HTT; all p_spin_ < 0.05) (Fig. 6E).

Finally, we asked whether subgroup-specific deviation maps preferentially localize to regions implicated in particular cognitive functions using Neurosynth meta-analytic maps (123 terms). For each subgroup, we correlated the average deviation map with probabilistic activation maps and corrected for multiple comparisons across terms (Fig. 6F). ASD1 showed selective positive associations with domains related to reasoning and executive control (‘Reasoning & Decision-Making’: *r* = 0.060, p_FDR_ = 0.037; ‘Executive/Cognitive Control’: *r* = 0.074, p_FDR_ = 0.007) and a negative association with perceptual systems (‘Perception’: *r* = − 0.078, p_FDR_ = 0.007). By contrast, ASD2 displayed a clear positive relationship with perceptual processing (‘Perception’: *r* = 0.086, p_FDR_ = 0.007), with no other categories surviving correction. These functional–anatomical dissociations dovetail with the molecular and hierarchical results: ASD1’s frontal/executive bias aligns with its frontal/motor loadings and focal neurotransmitter associations, whereas ASD2’s sensory/perceptual fingerprint matches its sensorimotor–visual embedding and broad receptor/transcriptomic coupling.

## 3 Discussion

In the present study, we combined subject-wise morphometric similarity (MIND) networks with HBR normative modeling to map individualized cortical atypicality in a large multisite sample of individuals with autism. Three main findings emerge. First, autism is associated with an increased prevalence of extreme, regionally focal morphometric deviations that are highly individualized and therefore poorly captured by conventional group averages. Second, these deviations are spatially structured. They tend to redistribute morphometric similarity away from association and heteromodal cortex toward sensory and visual territories and align with canonical cortical gradients and molecular maps. Third, individualized deviation maps cluster into reproducible subgroups that differ in hierarchical embedding, molecular coupling, and functional associations.

A persistent problem in structural neuroimaging of autism is the sheer inconsistency of findings. Some studies report regional hypertrophy and others atrophy[42, 43]. These discordant outcomes have fed a long-running debate about whether ASD is characterized by a coherent neuroanatomical signature or by heterogeneous, subject-specific alterations[44]. By building calibrated, per-region normative models and inspecting subject-wise deviation maps, we showed that small population-average effects can mask a more complex reality in which a minority of individuals carry large, focal departures from expected morphometry while many individuals show regional measures that overlap the normative range. Practically, this may suggest one major source of prior inconsistency, that is, group-level contrasts are highly sensitive to outliers and to the assumption of a single homogeneous effect.

Interestingly, the removal of outliers (individuals with highly atypical morphometric similarity) abolished the conventional case–control differences altogether (Fig. S9), demonstrating that those nominal group effects were driven largely by an atypical subset rather than a subtle, spatially consistent population shift. This pattern accords with other normative-modeling work that has emphasized single-subject heterogeneity in autism[45, 13, 15, 46], as well as other psychiatric and developmental cohorts [47, 48]. This further underscores that normative, uncertainty-aware approaches are essential for interpreting neuroanatomical variability in ASD.

Why deviations concentrate in particular cortical territories? We found that ASD group-level mean z-map showed reliable spatial structure rather than randomness. This might be that atypicality in ASD is constrained by the brain’s multiscale architecture[35, 22, 49, 29]. Region-wise mean deviations correlate negatively with the principal transcriptomic axis and with measures of receptor repertoire (receptor PC1 and receptor entropy), while showing positive associations with metabolic and myelin-rich maps. In other words, transmodal association territories that are transcriptomically distinct and carry diverse receptor portfolios tend to show reductions in morphometric similarity, whereas metabolically active, myelinated sensory territories more often show increased similarity.

This pattern is consistent with anatomical homophily, the idea that regions similar in microstructure, gene expression and chemoarchitecture also share connectivity and developmental trajectories[50]. This may suggest the vulnerability to morphometric reconfiguration in ASD is channeled by the cortex’s intrinsic molecular and metabolic landscape. This pattern invites specific causal hypotheses. For example, late-maturing association cortices may exhibit greater sensitivity to genetic or environmental perturbations[51]; conversely, sensory territories may manifest compensatory or activity-dependent increases in similarity tied to metabolic or myelination processes[52].

Importantly, morphometric deviation is not only a regional phenomenon. It also reconfigures interregional similarity. We found a large, asymmetric component concentrated in primary visual and higher-order visual nodes, dominated by increased between-region similarity in ASD but with focal reductions in selected association pairings (Fig. 3). Crucially, most significant changes were cross-network rather than within-network, indicating that ASD preferentially alters how different functional systems relate to one another rather than uniformly strengthening or weakening local cohesion. This selective reweighting is consistent with reports of altered functional-gradient maturation and atypical sensory–association transitions in ASD[53, 54, 55] and suggests that morphometric similarity reorganization may underlie or reflect altered integration/segregation balance in cortical processing[56, 57]. An open question is whether these structural-similarity changes precede functional gradient disruptions, follow them, or co-emerge with them during development.

Clustering individualized deviation maps uncovered two stable ASD subgroups with distinct hierarchical and molecular fingerprints. ASD1 carries net positive deviations concentrated in frontal and motor territories, exhibits weaker gene-gradient coupling, and is associated with a more circumscribed neurotransmitter profile (relatively greater dopaminergic weight). ASD2 is a spatially coherent, sensory-biased phenotype. Its deviations align toward the sensorimotor/visual pole, couple strongly to gene-expression and receptor gradients, and show a broad receptor fingerprint (notably serotonergic markers and several metabotropic/ionotropic systems). Furthermore, ASD1’s map correlates with executive and cognitive-control terms, while ASD2’s map aligns with perceptual and imagery-related terms. How might these phenotypes arise?

Regional developmental programs (timing of synaptogenesis[58], pruning[59], myelination [60], and receptor maturation[61]) create a spatially patterned landscape of vulnerability. Furthermore, perturbations (genetic risk, early-life environmental exposures, altered activity-dependent plasticity) interfacing with that landscape may yield locally heterogeneous outcomes[2]. For example, when perturbation aligns with molecular gradients it can produce a coherent, gradient-constrained phenotype (ASD2); when perturbation is more spatially focal or arises later it may produce a frontal/executive-biased phenotype (ASD1).

The cortical–molecular subgroups identified in our study generate testable hypotheses for stratified, mechanism-focused investigations[62]. ASD1’s fronto-motor profile suggests prioritizing circuit-targeted behavioural interventions or neuromodulatory strategies, whereas ASD2’s molecular coupling points to candidate pharmacological targets and to PET-based biomarker development. However, these imaging-derived subgroups must be validated against behavioural and clinical measures[63]; they are best understood as mechanistic hypotheses rather than clinical diagnoses. Future studies should determine whether subgroup membership predicts sensory sensitivity, executive function, treatment response, or developmental trajectory beyond conventional clinical stratifiers.

Several limitations deserve mention. The study is cross-sectional, thus normative z-scores describe deviation at single time points and cannot resolve developmental trajectories, and longitudinal hierarchical models are needed to distinguish delayed from divergent maturation. Our sample is multisite and not population representative; although HBR models explicitly account for site effects and we ran sensitivity checks, residual sampling and acquisition biases may remain, and much of the ABIDE data were collected in Western populations, which may limit generalizability of our results. Analytic choices such as parcellation scheme, morphometric feature set and clustering algorithm may also influence outcomes. Molecular annotations such as AHBA transcriptomics and PET-derived neurotransmitter and transporter maps are spatial proxies assembled from small or aggregated cohorts and do not measure subject-specific neurochemistry, so the spatial correlations we report generate candidate mechanisms rather than prove causality. Motion, medication, comorbidity and ascertainment differences vary across sites and were not uniformly controlled, and these factors can influence morphometric estimates. We modeled sex as a covariate but did not perform fully powered sex-stratified normative analyses or formal tests of sex-by-diagnosis interactions given the sex imbalance typical of ASD cohorts, so dedicated work is required to evaluate whether the deviation patterns we observe differ by sex.

In summary, rather than a single “ASD brain,” we show the cortex exhibits focal, gradient-constrained, and molecularly tethered reconfigurations that are concentrated in a minority of individuals and that resolve into reproducible cortical–molecular phenotypes. Normative, subject-wise modeling of architectonically informed structural-similarity networks provides an important path for illuminating this heterogeneity and for generating mechanistic hypotheses. Embracing heterogeneity as signal rather than noise will be essential if neuroimaging is to advance mechanistic understanding and enable stratified interventions in ASD.

## 4 Methods

### 4.1 Participants

The study included participants from both TD and ASD cohorts. Structural MRI (sMRI) data was initially obtained from Autism Brain Imaging Data Exchange (ABIDE) I (17 sites, ASD/TD: 539/573, 7-64 years) and II (19 sites, ASD/TD: 487/557, 5-64 years)[64, 65]. All data were fully anonymized in accordance with HIPAA. Diagnostic criteria, acquisition parameters, informed consent procedures, and site protocols are available from the ABIDE consortium (https://fcon_1000.projects.nitrc.org/indi/abide/). The initial dataset was screened using pre-specified inclusion criteria: (i) nearly full-brain coverage and visually acceptable image quality on native T1 images, and (ii) full-scale IQ > 70. After subject-level screening and the Quality control (QC) pipeline described below, sites with fewer than 10 participants remaining in either the ASD or TD group were removed from further analysis. The final sample consisted of 1344 individuals (ABIDE I: 656 participants from 13 sites; ABIDE II: 688 participants from 10 sites) (Table S1; Fig. S1).

### 4.2 Neuroimaging data acquisition and preprocessing

T1-weighted images were processed with FreeSurfer v7.4.1[66, 67]using the standard recon-all pipeline with default parameters. The pipeline steps included motion correction, non-uniform intensity (bias) normalization, skull stripping, affine registration to MNI305 space, segmentation of subcortical white and deep gray matter, cortical white/pial surface reconstruction, automatic topology correction, surface inflation, and spherical registration to an average atlas to establish inter-subject cortical correspondence.

QC was implemented to ensure the integrity of raw images and surface reconstructions. First, raw image quality was quantified on the native T1 using the CAT12 Image Quality Rating (IQR); datasets with IQR ≤ 70 were excluded (threshold chosen to remove scans with substantial artifacts or poor contrast). Second, FreeSurfer recon-all was executed for retained subjects and Euler numbers were computed. Subjects with Euler numbers more negative than − 120 (i.e., Euler < − 120, corresponding to ≥ 2.6 median absolute deviations below the sample median) were excluded as topological outliers. Third, reconstructed cortical and subcortical surfaces were visually inspected by experienced raters to identify residual reconstruction failures (for example, skull-strip errors or gross pial/white segmentation errors); reconstructions judged irreparable were excluded. Site-level exclusions (sites with <10 participants in either group) were applied after subject-level screening and QC. The distribution of Euler numbers across hemispheres for the retained sample is shown in Fig. S2.

### 4.3 MIND network construction

For each subject, we defined 308 cortical regions by subdividing the Desikan–Killiany atlas into equal-area parcels (~ 500 mm^2^ each)[68, 69]. Within each parcel, we extracted the distribution of five vertex-wise morphometric features: cortical thickness, surface area, gray matter volume, mean curvature, and sulcal depth. We then computed a pairwise similarity between regions *a* and *b* as the symmetrized Kullback–Leibler (KL) divergence between their multivariate feature distributions[33]. Using a k-nearest-neighbor estimator for multivariate divergence, we convert divergence to a bounded similarity:

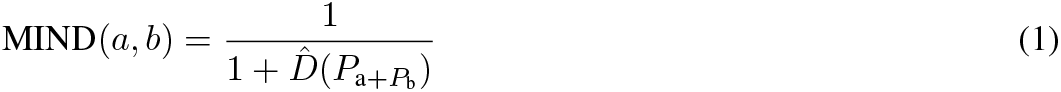

where 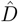 is the estimated symmetric KL divergence. Each subject’s result is a 308×308 MIND similarity matrix. All feature vectors were standardized (z-scored) across the brain prior to MIND computation to ensure comparability, and vertices with zero values in area/volume were excluded to avoid unfeasible inputs.

### 4.4 Hierarchical Bayesian regression normative modeling

We fit a HBR model separately for each ROI’s mean MIND value (weighted degree / hubness of each regional node) across the cohort, using the PCNtoolkit framework[17]. The model predicts the expected mean MIND value as a function of age (modeled with a B-spline basis), sex (linear term), and includes scanner site as a random-effect (random intercepts and slopes) to capture multi-site variance.

Specifically, we parameterized the expected mean *µ* and variance *σ*^2^ of the ROI’s mean MIND value using a B-spline basis on age (to capture non-linear developmental trends), a linear term for sex, and an intercept. Site was encoded as a categorical “batch” variable, and we allowed both random intercepts and random slopes by site for *µ* (and similarly for *σ*) in the hierarchical model. This Bayesian partial-pooling approach “borrows strength” across sites while allowing site-specific adjustments, controlling for potential scanner effects without sacrificing generalizability.

We trained the HBR models on an age- and sex-matched subset of the typically developing (TD) cohort (80% of TD subjects, with stratification by site) and tested on the held-out 20% of TD plus all ASD cases. From the posterior predictive distributions, we extracted for each test subject and each ROI a z-score:

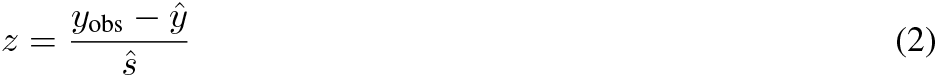

where *y*_obs_ is the observed ROI MIND value, *ŷ* the posterior mean prediction, and *ŝ* the predictive standard deviation. By definition, under a correctly specified normal model these z-scores would follow a unit normal distribution for the TD reference group, while ASD subjects may show outlier deviations. Model performance was quantified via standard normative metrics, including explained variance (EXPV), standardized mean squared error (SMSE), and mean standardized log-loss (MSLL) among others.

### 4.5 Network-based statistic

We tested for topologically coherent differences in interregional morphometric similarity with the NBS[70]. For each subject we first residualized the 308×308 MIND similarity matrix to remove linear effects of age, sex, site and the age×sex interaction; edge values were entered into edge-wise two-sample t tests (ASD vs TD). A primary ≥ t-statistic threshold of 3.2 was applied to identify suprathreshold edges, and connected components (sets of nodes joined by suprathreshold edges) were formed separately for positive and negative contrasts. Statistical inference used nonparametric permutation of group labels (5,000 permutations) to construct the null distribution of maximal component extent; component-wise p values were computed as the proportion of permutations with a maximal component size ≥ the observed.

### 4.6 Identifying distinct morphometric similarity subgroups

HYDRA is a semi-supervised method that employs a supervised machine learning algorithm to determine boundaries that separate controls from patients while simultaneously identifying patient-specific subgroups [71]. Reproducibility of the subgroup solution is assessed by using an internal cross-validation cycle to identify subgroups. Key advantages of HYDRA, over other machine learning techniques, are that it disposes of the need for a priori specification of the number of clusters and does not use similarity measures for clustering as such measures are susceptible to the effect of non-specific factors such as age and sex. The detailed code can be found on https://github.com/evarol/HYDRA.

Classification in HYDRA is based on indices of deviation between a clinical and the healthy reference group; healthy individuals are separated from the clinical sample using a convex polytope formed by combining multiple linear hyperplanes. This confers an additional advantage to HYDRA because the multiple hyperplanes model potential heterogeneity within clinical samples while their combination extends linear max-margin classifiers to the non-linear space. The regional z scores of all ASD and TD individual entered into the algorithm as input features and age and sex were modeled as covariates. We used 5-fold cross validation to determine the clustering stability. The resultant clustering solutions were evaluated using the adjusted Rand index (ARI), adjusted for the chance grouping of elements; the solution with the highest ARI was chosen.

### 4.7 Biologically annotated brain maps

To investigate whether the observed deviation patterns are associated with underlying molecular architecture, we systematically compared cortical maps of mean MIND deviations to multiple micro-architectural annotated brain maps. These measures included gene expression, neurotransmitter receptor densities, excitatory-inhibitory ratio, glucose metabolism, myelination, synapse density, and receptor entropy.

Gene expression data was obtained from the Allen Human Brain Atlas (AHBA)[72] and processed using the abagen toolbox[73]. Expression data was extracted for the 308 parcellation using bidirectional left-right mirroring and missing values were filled using centroid coordinates. A total of 10,281 genes with differential stability greater than 0.1 were retained. The resulting region-by-gene matrix (308 regions × 10,281 genes) was averaged across donors and missing values were imputed using k-nearest neighbors (k=5). The first principal component was computed after z-scoring the imputed expression data, representing the primary axis of gene expression variation across cortical regions.

Receptor densities were collected for 19 different neurotransmitter receptors and transporters across 9 neurotransmitter systems from 25 different PET tracer studies as described in Hansen et al[74]. These include dopamine (D1[75], D2[76, 77], DAT[78]), norepinephrine (NET[79]), serotonin (5HT1a[80], 5HT1b[80, 81], 5HT2a[82],5HT4[82], 5HT6[83], 5HTT[82]), acetylcholine (*α*4*β*2[84], M1[85], VAChT[29, 86, 87]), glutamate (mGluR5[29, 88, 89], NMDA[90]), GABA (GABAa[91]), histamine (H3[92], cannabinoid (CB1[93]), and opioid (MOR[47]).

Individual PET images were sampled along the cortical ribbon at a fractional distance of 0.4 to 0.6 to minimize partial volume effects, and warped to MNI-ICBM 152 space using FreeSurfer’s CVS registration parameters. Receptors with multiple studies were combined using sample-size weighted averages after z-scoring individual datasets. The receptor gradient was computed as the first principal component of the parcellated receptor density matrix, representing the primary axis of variation in receptor organization across the cortex. Additionally, an excitatory/inhibitory ratio was calculated as the mean density of excitatory receptors (5HT2a, 5HT4, 5HT6, D1, mGluR5, *α*4*β*2, M1) divided by the mean density of inhibitory receptors (5HT1a, 5HT1b, CB1, D2, GABAa, H3, MOR). The receptor entropy was computed as the Shannon entropy of normalized receptor densities across all 19 receptors for each region.

Cortical glucose metabolism was derived from 18F-FDG PET data from 33 healthy adults (19 female, mean age 25.4 ± 2.6 years) as described in Vaishnavi et al[94]. Myelination measures were obtained from T1w/T2w ratios from the Human Connectome Project (S1200 release)[95, 96] for 417 participants (age 22-37 years, 193 males) acquired on a Siemens Skyra 3T scanner at 0.7 mm isotropic resolution, with gradient and bias field corrections applied[74].

Synapse density was measured in 76 healthy adults (45 males, mean age 48.9 ± 18.4 years) using 11C UCB-J PET tracer targeting synaptic vesicle glycoprotein 2A (SV2A), with 90-minute dynamic scans and SRTM2 modeling using centrum semiovale as reference[74]. All datasets were parcellated to 308 cortical regions and z-scored for consistent comparison across biological measures.

### 4.8 Partial least squares regression analysis

To investigate the molecular basis of ASD subgroup differences, we applied PLS regression to identify relationships between regional gene expression profiles and MIND deviation patterns[97, 98]. Analyses were conducted at the region level using a gene matrix of 10,281 genes. For each analysis the predictor matrix X consisted of regional gene-expression vectors (regions × genes) with each gene column z-scored across regions to remove gene-specgific mean and scale differences. The response vector Y was the subtype-specific mean MIND deviation per region (the normative z-scores derived from the hierarchical Bayesian regression). PLS regression decomposes the cross-covariance between X and Y into orthogonal LVs, each formed by a pair of gene and brain weighting vectors that capture maximally covarying multivariate patterns. We fitted ten components and focused inference on the first LV while reporting the variance explained across the retained components.

Statistical significance of LVs was assessed with nonparametric permutation testing. For per-LV tests we generated null distributions by permuting the regional order of Y (1,000 permutations), refitting PLS regression to each permuted dataset, and estimating the probability of observing an LV correlation at least as large as the empirical value. We focused on the first LV and on its regional brain scores and gene loadings, which together identify the spatial gene expression profile most tightly coupled to the subgroup MIND deviation pattern.

To assess the stability of PLS regression weights we performed bootstrap resampling across regions (1,000 bootstrap samples). Bootstrap standard errors of gene and regional weights were used to compute bootstrap ratios (weight / bootstrap SE), which we interpret as indices of weight reliability across resamples. To evaluate out-of-sample generalizability of our result, we ran 1,000 randomized cross-validation splits. Rather than randomly splitting brain regions into training and testing sets, we selected the 75% of regions closest in Euclidean distance to a randomly chosen source node as the training set, with the remaining 25% as the testing set. This approach ensures that spatially adjacent regions are grouped together, preventing data leakage that could occur when nearby regions with similar values are split across training and testing sets. For each split we estimated PLS weights on the training regions and projected the held-out regions to compute test correlations between gene and brain LV scores. Thus, the distribution of test correlations characterizes expected generalization and the training–test gap.

### 4.9 Dominance analysis

We used dominance analysis to quantify the relative contribution of seven biological attributes to the regional pattern of MIND deviation. Dominance analysis decomposes the total explained variance of a multiple linear regression into additive contributions for each predictor by fitting every possible model subset[99]. For a model with p predictors, we computed the change in adjusted R^2^ produced by adding each predictor to all possible submodels and averaged those increases to obtain the predictor’s total dominance. The sum of all predictor dominances equals the adjusted R^2^ of the full model, which provides an intuitive partitioning of model fit across predictors.

We modeled mean regional deviation patterns using biological attributes, as well as using neurotransmitter receptor and transporter density maps. Analyses were performed separately for the overall ASD map and for each ASD subtype (ASD1 and ASD2). Adjusted R^2^ for each empirical model was computed from least squares fits.

Because cortical maps are spatially autocorrelated, we assessed spatial generalizability using a distance-dependent cross-validation[100]. For each region in turn, we defined the training set as the 75% of regions closest in Euclidean space and the test set as the remaining 25%. Models were fit on the training set and predictions evaluated in the test set. For each held-out fold we stored the Pearson correlation between predicted and observed regional deviations; the distribution of these correlations summarizes out-of-sample spatial generalization. Dominance rankings were compared across groups by correlating dominance vectors.

### 4.10 Neurosynth functional annotation

To investigate the functional correlates of brain structural deviations in ASD subgroups, we utilized the Neurosynth meta-analytic database[101]. This comprehensive resource integrates findings from over 15,000 fMRI studies, providing probabilistic activation maps that quantify the association between specific cognitive terms and regional brain activity. These maps offer a quantitative representation of how regional brain activity relates to various psychological processes. Our analysis focused on 123 cognitive and behavioral terms from the Cognitive Atlas[101], ranging from broad categories (e.g., *attention, emotion*) to specific cognitive processes, behaviors, and emotional states. These terms were systematically organized into 11 functional categories, including: *Action, Learning and Memory, Emotion, Attention, Reasoning and Decision Making, Executive/Cognitive Control, Social Function, Perception, Motivation, Language*, and *Other*.

To assess whether terms within each functional category were significantly correlated with brain deviation patterns in each ASD subgroup, we employed a two-sided permutation test. This involved randomly permuting the category assignments for each term 10,000 times and subsequently computing the average correlations from these permuted categories to generate a null distribution. Empirical correlations were then compared against this null distribution to derive p-values. Finally, these p-values were corrected for multiple comparisons using the Benjamini-Yekutieli false discovery rate procedure.

### 4.11 Spatial autocorrelation-preserving null annotations

To address the potential bias in statistical significance assessment due to spatial autocorrelation in brain imaging data[102, 103], we implemented a complementary approach for generating null distributions that preserve the spatial structure of cortical maps. We employed spin permutation tests[102]. This method generates spatially autocorrelated null maps by randomly rotating the spherical projection of cortical surface data while preserving spatial contiguity and hemispheric symmetry. For each correlation analysis between brain maps, we compared the empirical Pearson correlation coefficient to a null distribution of 10,000 correlations between the empirical map and the randomly rotated projections of the comparison map, computing p-values as the proportion of null correlations with absolute values greater than or equal to the observed correlation.

## 5 Data availability

All neuroimaging data is openly available through the ABIDE consortium: https://fcon_1000.projects.nitrc.org/indi/abide/; the Allen Human Brain Atlas is available at https://human.brain-map.org; the neuromaps toolbox, utilized for brain mapping analyses, is available at https://github.com/netneurolab/neuromaps.

## 6 Acknowledgements

We thank all study participants and investigators contributing to the ABIDE consortium.

---

**Table S1:**
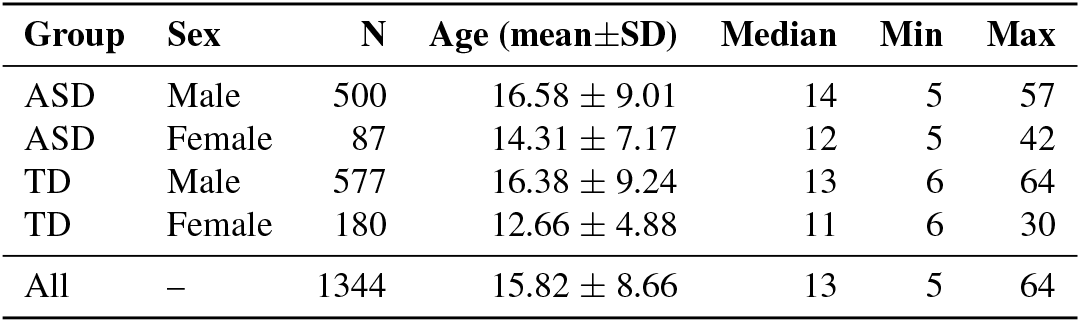
Participant demographics by group and sex.

**Figure S1.**
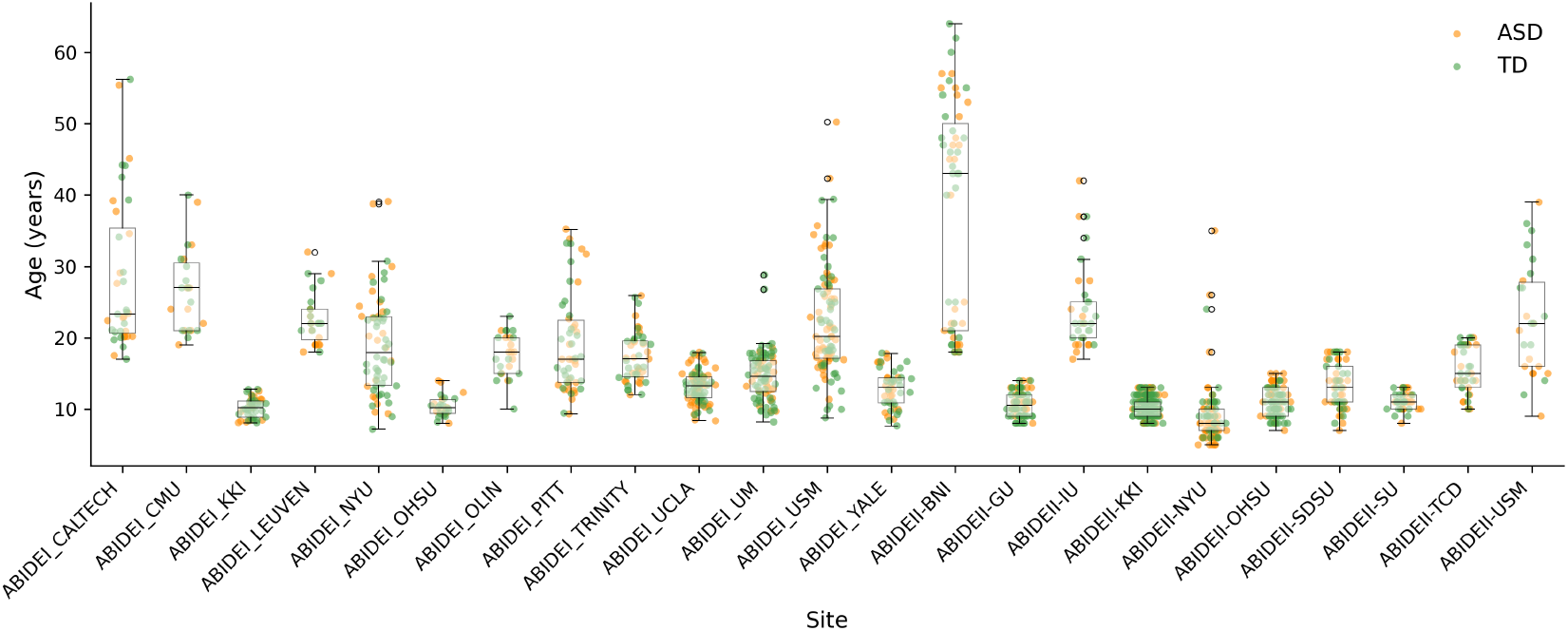
Participant age distribution by site. Site-wise age distribution for 23 ABIDE sites; ASD (n = 587) in orange and TD (n = 757) in green.

**Figure S2.**
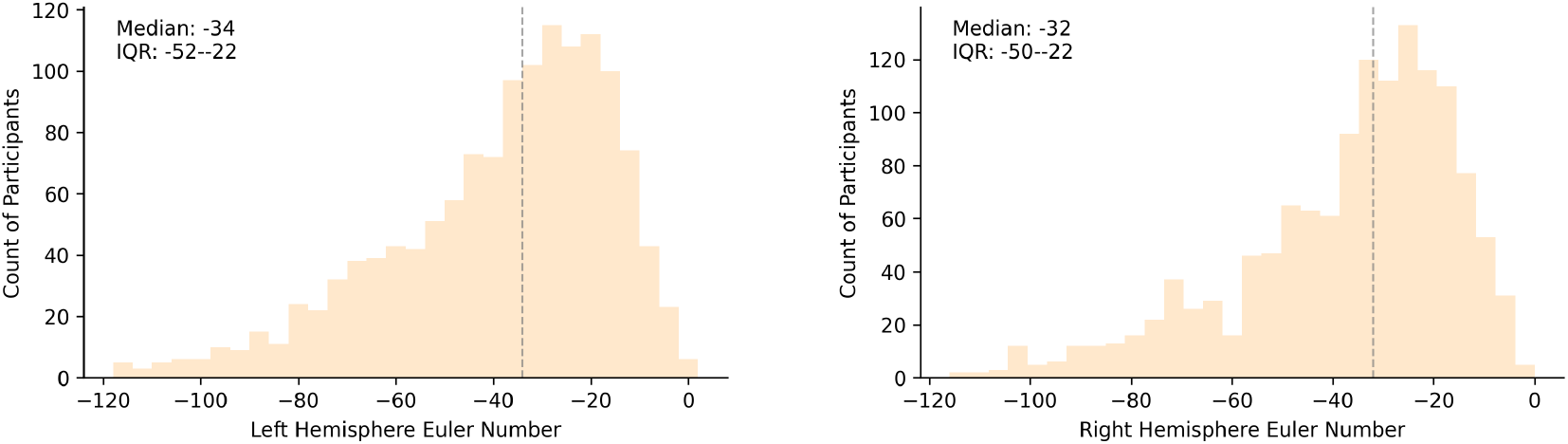
Distribution of cortical Euler numbers. Histograms of Euler numbers by hemisphere across participants; medians and IQRs annotated. Vertical dashed lines indicate the median values (left: −34, right: −32), with interquartile ranges displayed in each panel (left: −52 to −22, right: −50 to −22). The distributions reveal that both hemispheres show similar patterns of cortical complexity, with left hemisphere showing slightly greater folding (mean: −39.6±23.0) compared to right hemisphere (mean: −37.1±21.9). Euler numbers ranged from −118 to 2 across both hemispheres.

**Figure S3.**
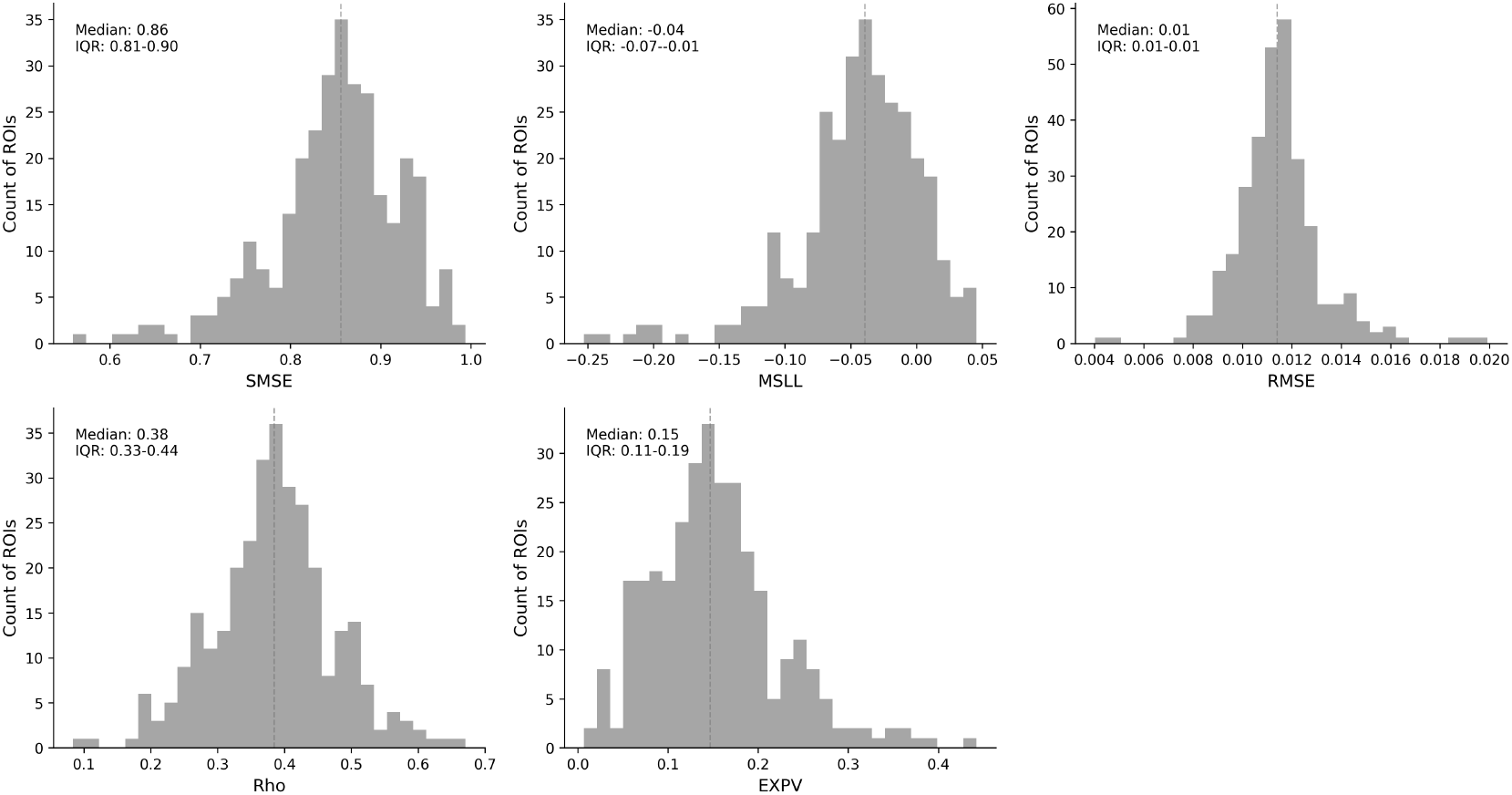
Per-ROI HBR performance metrics. Histograms of standardized mean squared error (SMSE), mean standardized log loss (MSLL), root mean squared error (RMSE), correlation coefficient (Rho) and explained variance (EXPV) across 308 ROIs demonstrating overall model adequacy for normative prediction.

**Figure S4.**
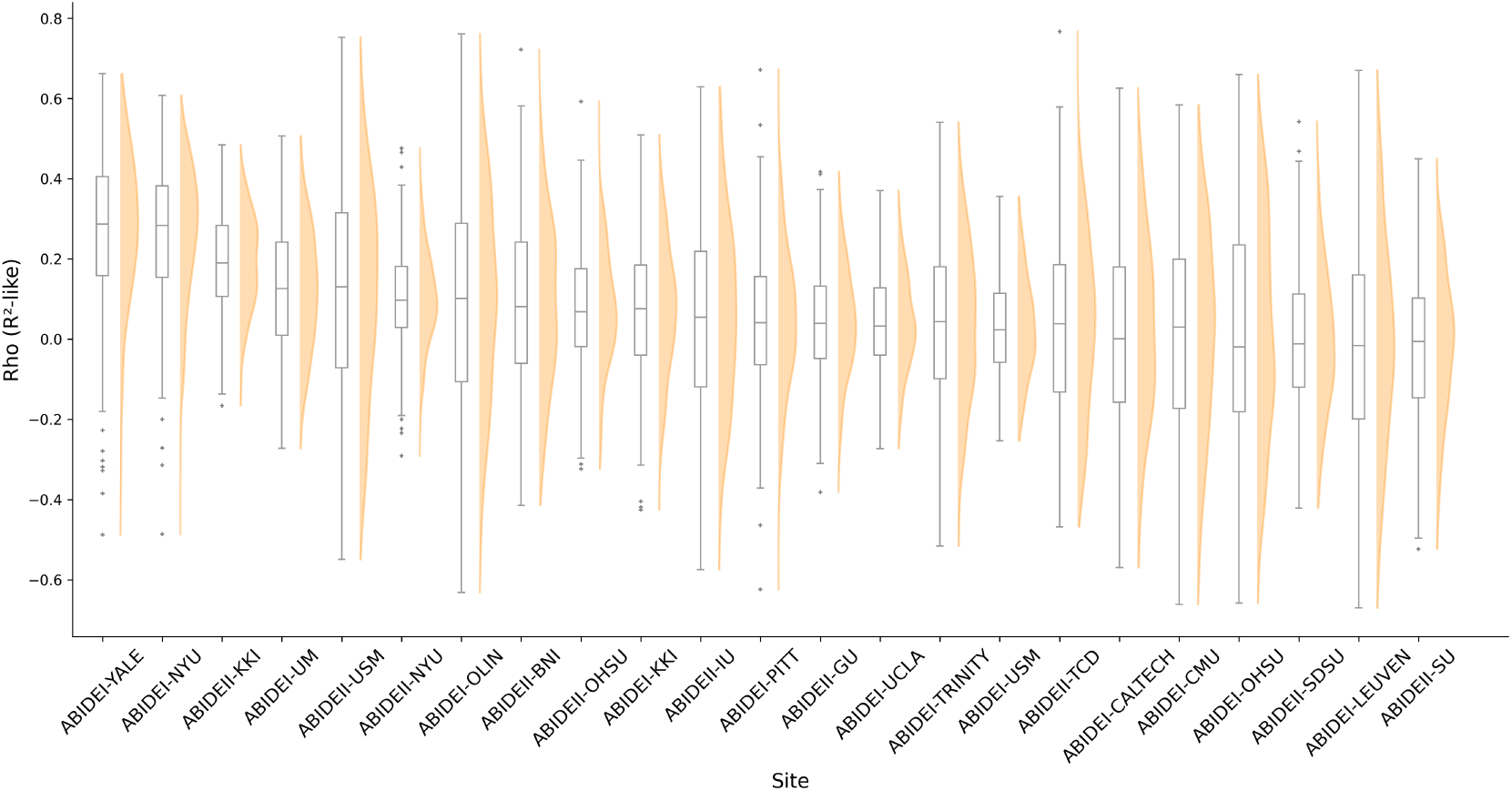
Site-level normative model performance. Site-wise Rho distributions (box + density) illustrating between-site heterogeneity in model fit.

**Figure S5.**
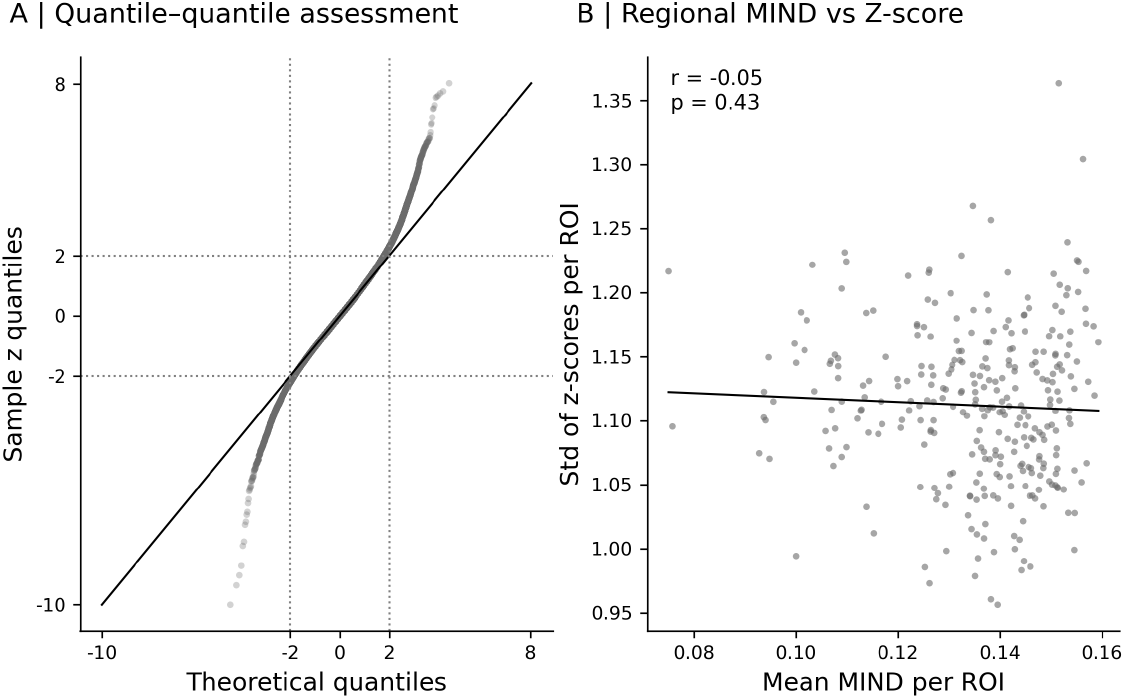
Z-score calibration and regional dispersion. (A) Q–Q plot of empirical z-scores vs theoretical Gaussian; tails indicate extreme outlier prevalence. (B) Lack of association between mean regional MIND and z-score SD (r = −0.05, p = 0.43).

**Figure S6.**
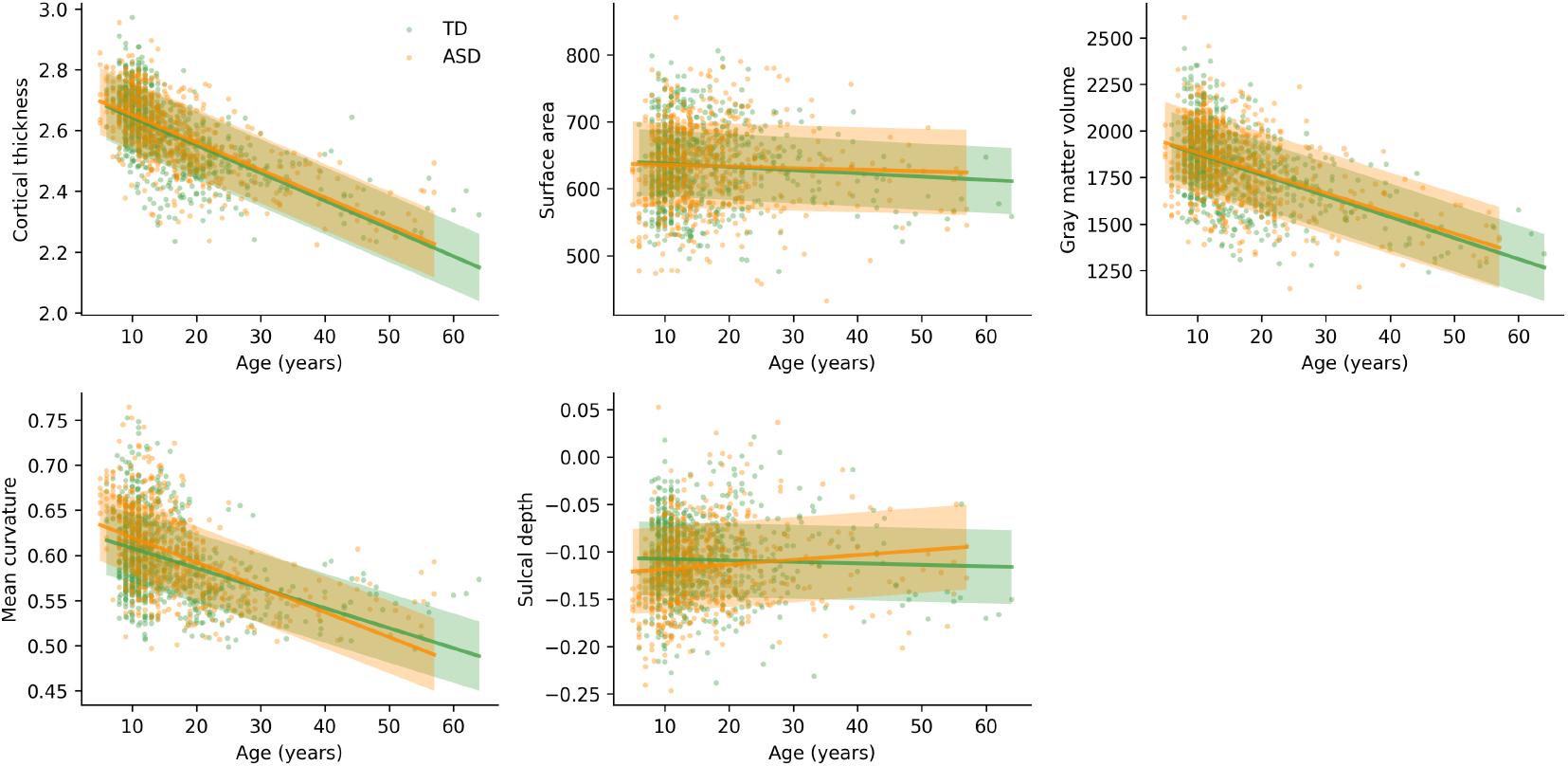
Developmental trajectories of morphometric features. Region-averaged age trends for cortical thickness, surface area, gray matter volume, mean curvature and sulcal depth across ASD and TD participants.

**Figure S7.**
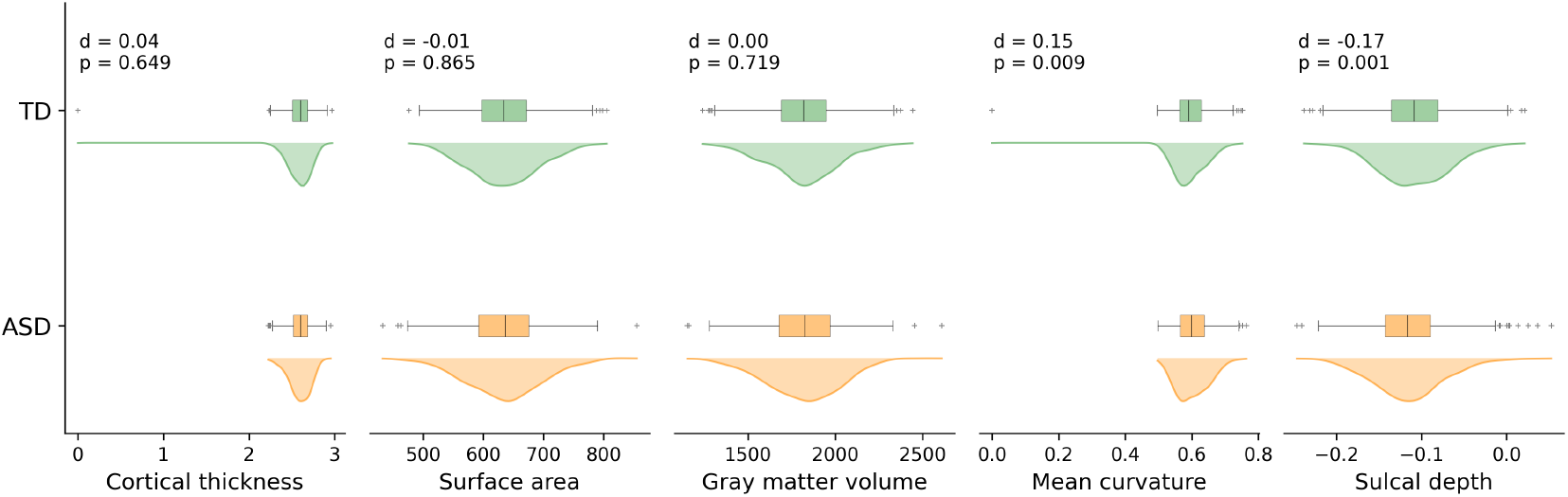
Group comparisons of morphometric features. ROI-averaged comparisons for five morphometric measures. Statistical comparisons were performed using non-parametric tests (Mann-Whitney U) to account for non-normal distributions. The analysis reveals group differences in cortical morphometry, with effect sizes calculated using Cohen’s d for continuous variables. Data are presented as box plots with overlaid kernel density estimation (KDE) distributions to show the shape of the data distribution.

**Figure S8.**
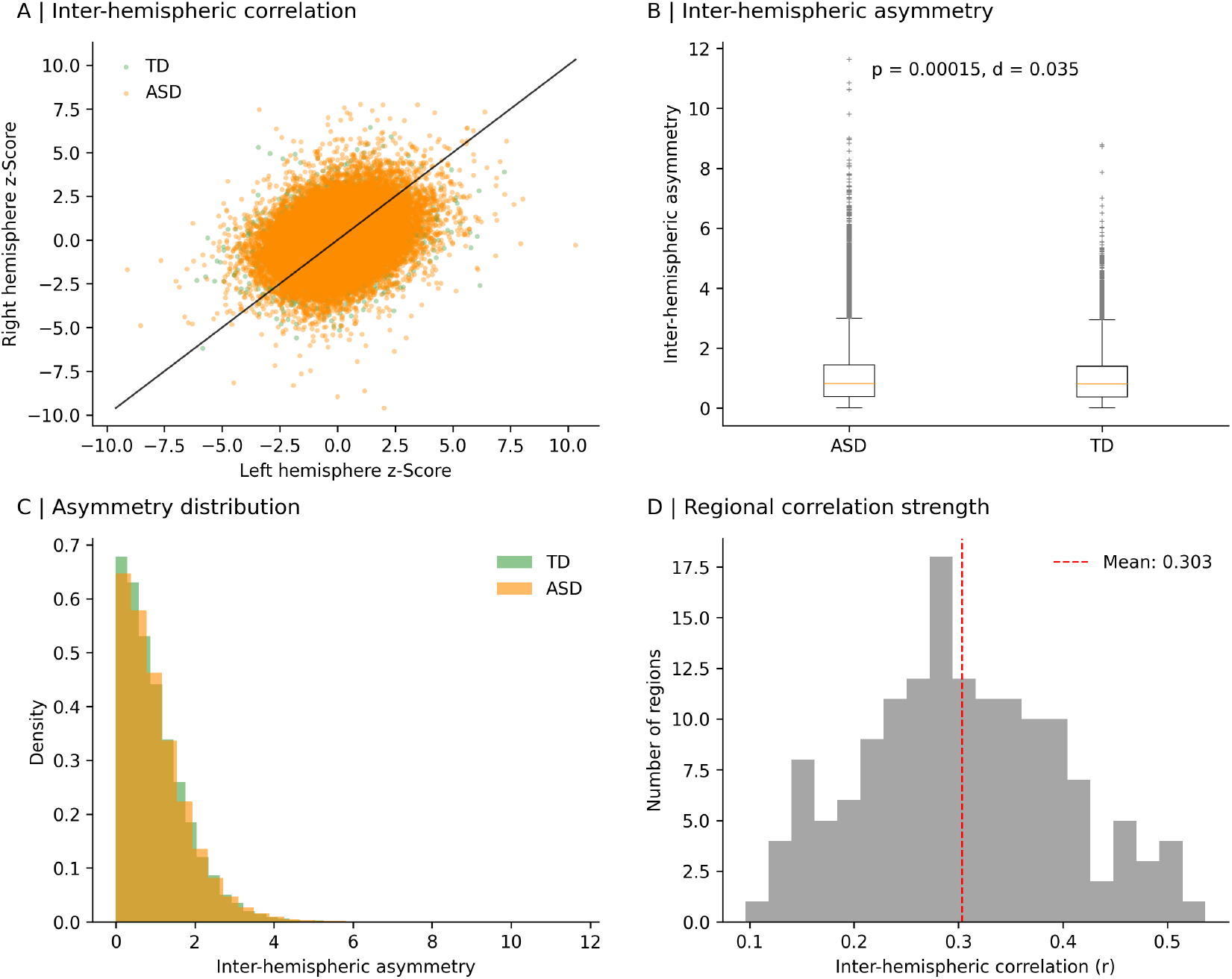
Inter-hemispheric correspondence and asymmetry. (A) Scatter plot shows the relationship between left and right hemisphere z-scores for homologous regions, with each point representing a participant’s z-score pair for a given region and the diagonal line indicating perfect correlation. Both ASD and TD groups show strong positive correlations (ASD: r = 0.311, p < 0.001; TD: r = 0.272, p < 0.001), indicating preserved inter-hemispheric structural symmetry in both groups. (B) Box plots comparing the magnitude of inter-hemispheric asymmetry (absolute difference between left and right z-scores) between ASD and TD groups, calculated as |left z-score - right z-score| for each homologous region pair, with the groups showing minimal differences (Cohen’s d = 0.035), suggesting preserved bilateral organization in ASD. (C) Histogram showing the distribution of inter-hemispheric asymmetry values for ASD and TD groups, with both distributions being similar and most values clustered near zero, indicating that extreme inter-hemispheric differences are rare in both groups. (D) The distribution of inter-hemispheric correlation coefficients across all homologous region pairs demonstrating that most regions exhibit moderate to strong inter-hemispheric correlations with few regions showing weak correlations.

**Figure S9.**
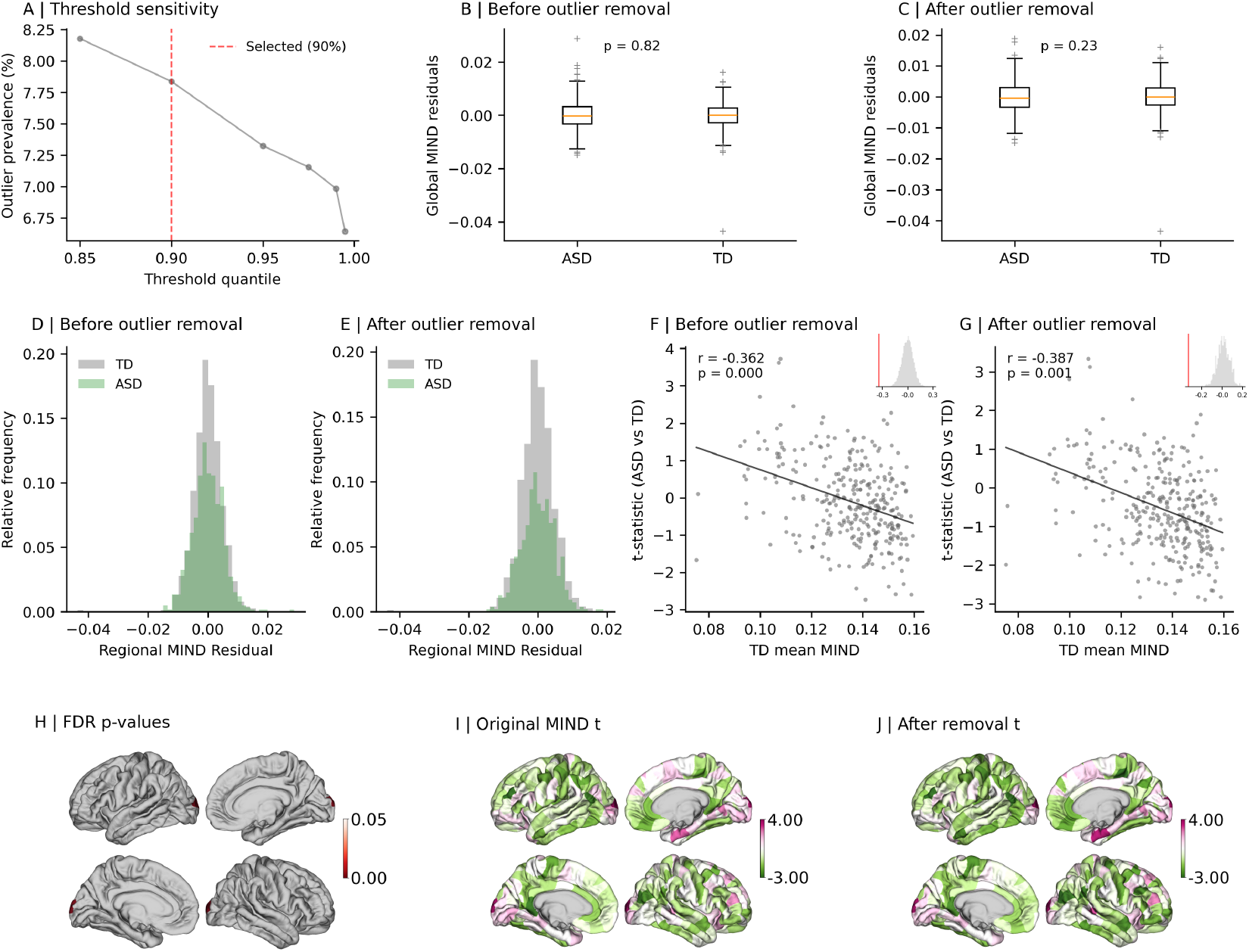
Outlier influence on conventional case–control differences. (A) Mahalanobis-distance-based outlier identification. The selected 90% threshold (red dashed line) identifies 46 ASD participants (7.8% of ASD sample) as statistical outliers. Box plots showing the comparison of global mean MIND residuals from linear regression models (controlling for age, sex, age × sex interaction, and site) for typically developing and ASD participants before (B) and after (C) outlier removal (Mann-Whitney U). Histograms showing the distribution of regional MIND residuals across all brain regions for TD and ASD participants before (D) and after (E) outlier removal. Scatter plot showing the relationship between mean TD MIND values and t-statistics from ASD vs. TD comparisons across brain regions before (F) and after (G) outlier removal. The correlation coefficient (r) and significance (p) from spatial permutation testing (spin test) are displayed. The inset histogram shows the null distribution from 10,000 spatial permutations, with the empirical correlation marked by a red line. Brain plot (H) shows FDR-corrected p-values for regions that survived multiple testing correction (p < 0.05) (2 out of 308 regions). After outlier removal no regions remained significant after FDR correction. Brain plots showing t-statistics from the case-control analysis comparing ASD to TD participants before (I) and after (J) outlier removal.

**Figure S10.**
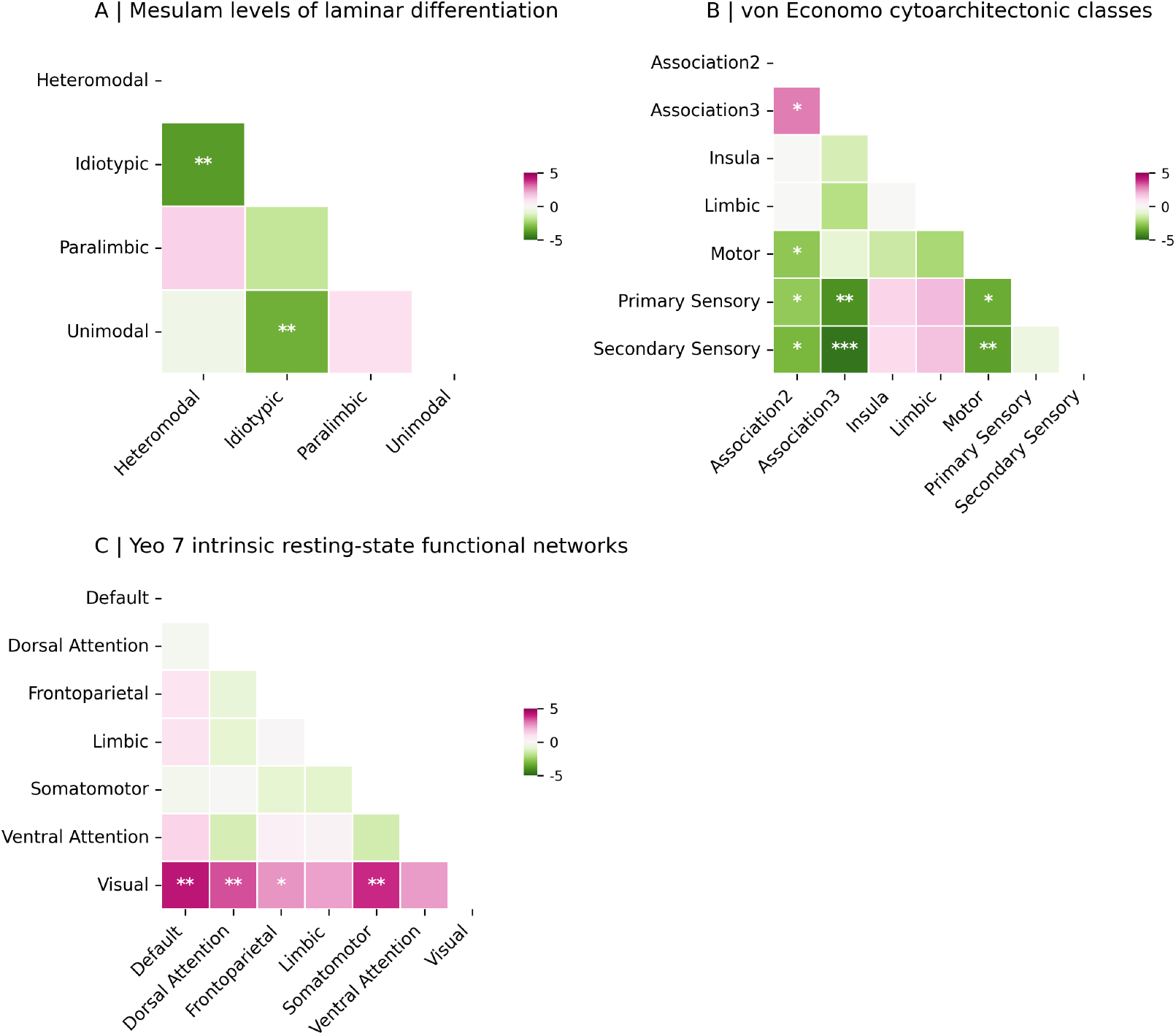
Between-zone contrasts across parcellation schemes. Heatmaps of pairwise zone comparisons for Mesulam, von Economo and Yeo7 partitions with permutation-based significance. Statistical significance is indicated by asterisks (***pFDR < 0.001, **pFDR < 0.01, * pFDR < 0.05).

**Figure S11.**
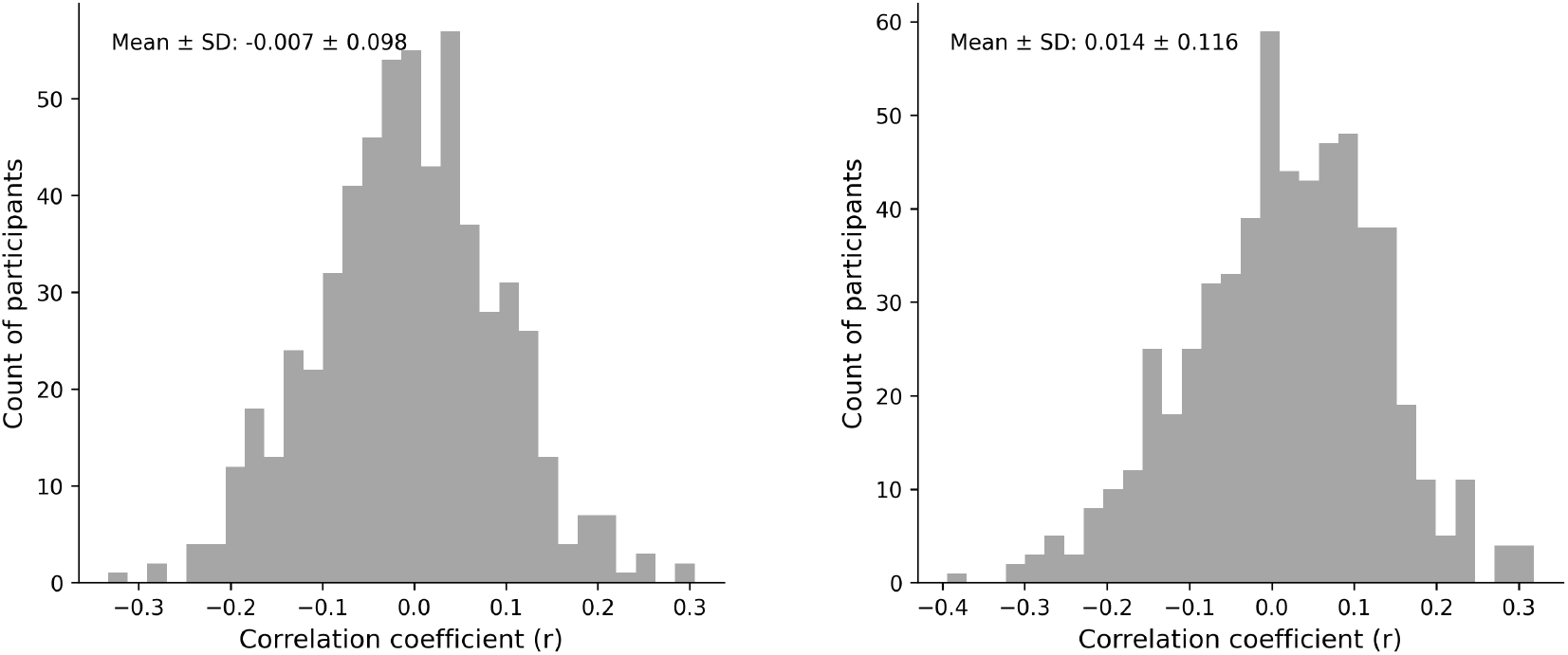
Subject-level alignment between canonical functional gradients and individual ASD morphometric deviations. Histograms of individual correlations between subject z-maps and canonical gradients (G1/G2).

**Figure S12.**
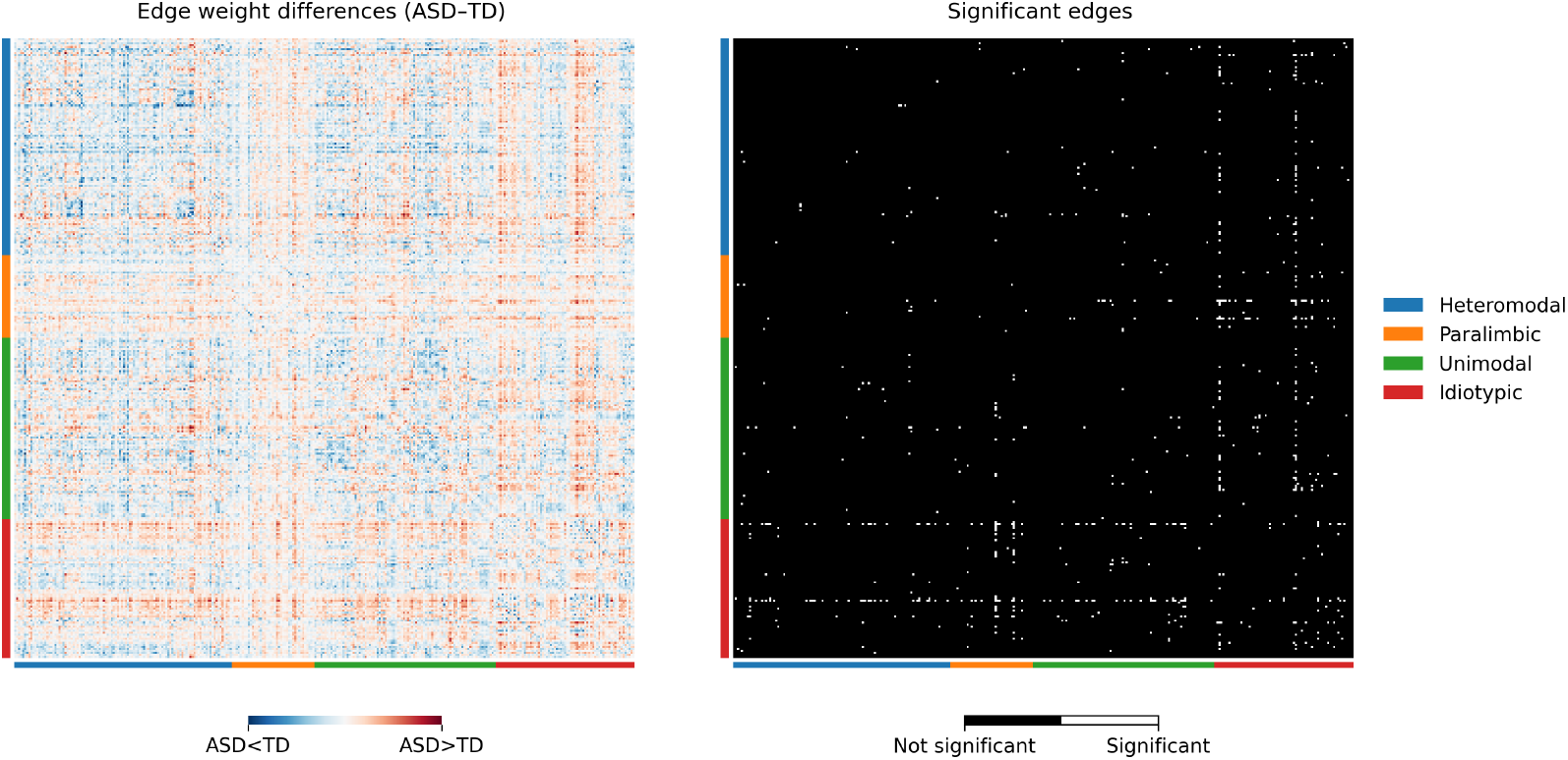
Edgewise contrasts and NBS-identified edges. (A) Unthresholded matrix of ASD – TD residualized MIND edge weights (308 × 308). Warm colors indicate stronger morphometric similarity in ASD; cool colors indicate stronger similarity in TD. Rows/columns are ordered by Mesulam zone (heteromodal, paralimbic, unimodal, idiotypic). (B) Binary map of edges surviving the NBS (primary t = 3.2; 5,000 permutations; component FWE p < 0.05). White pixels mark significant edges from the FWE-corrected component; black pixels indicate non-significant edges.

**Figure S13.**
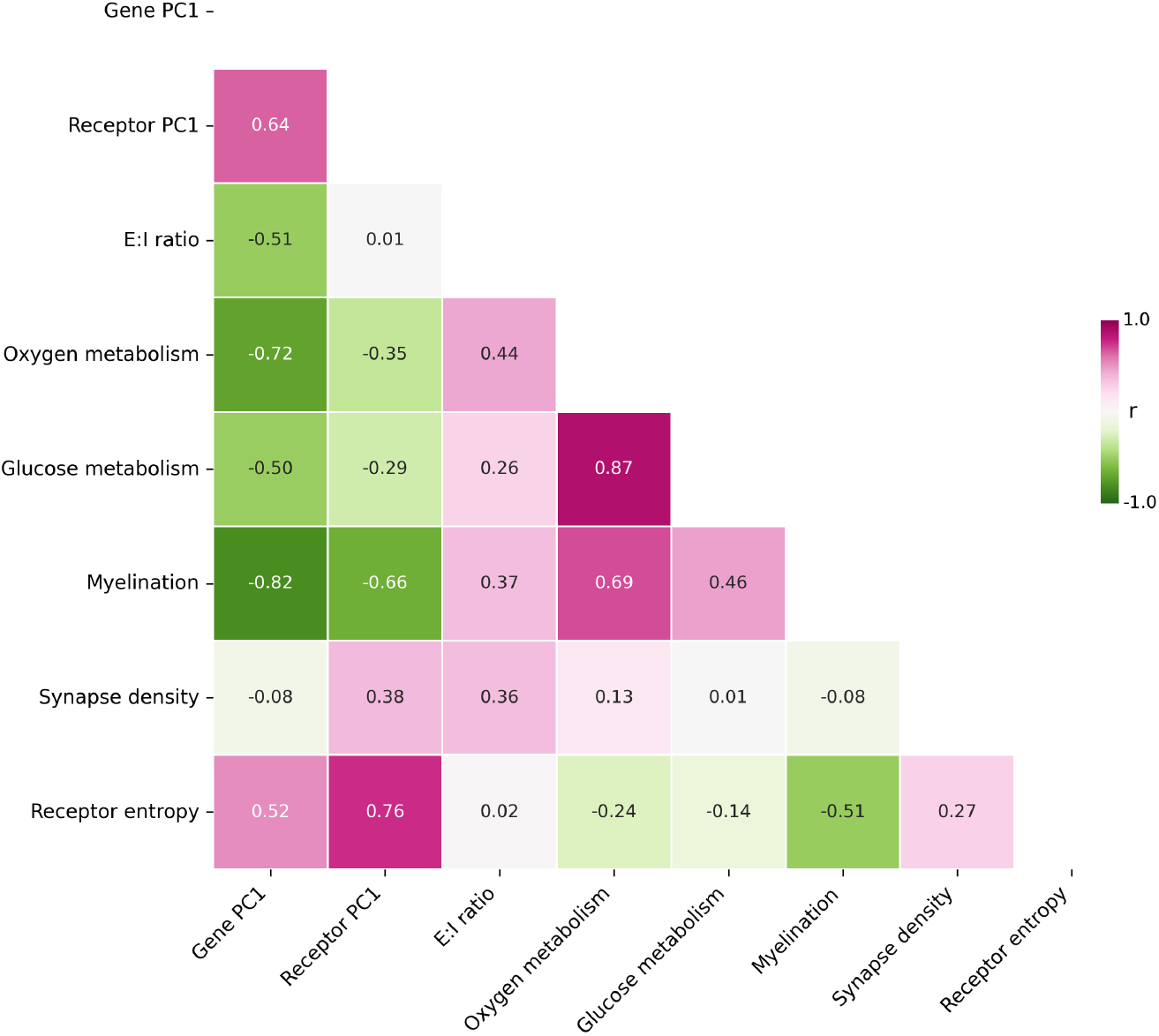
Pairwise correlations among biological attribute maps. Pearson correlation matrix across the set of biological maps.

**Figure S14.**
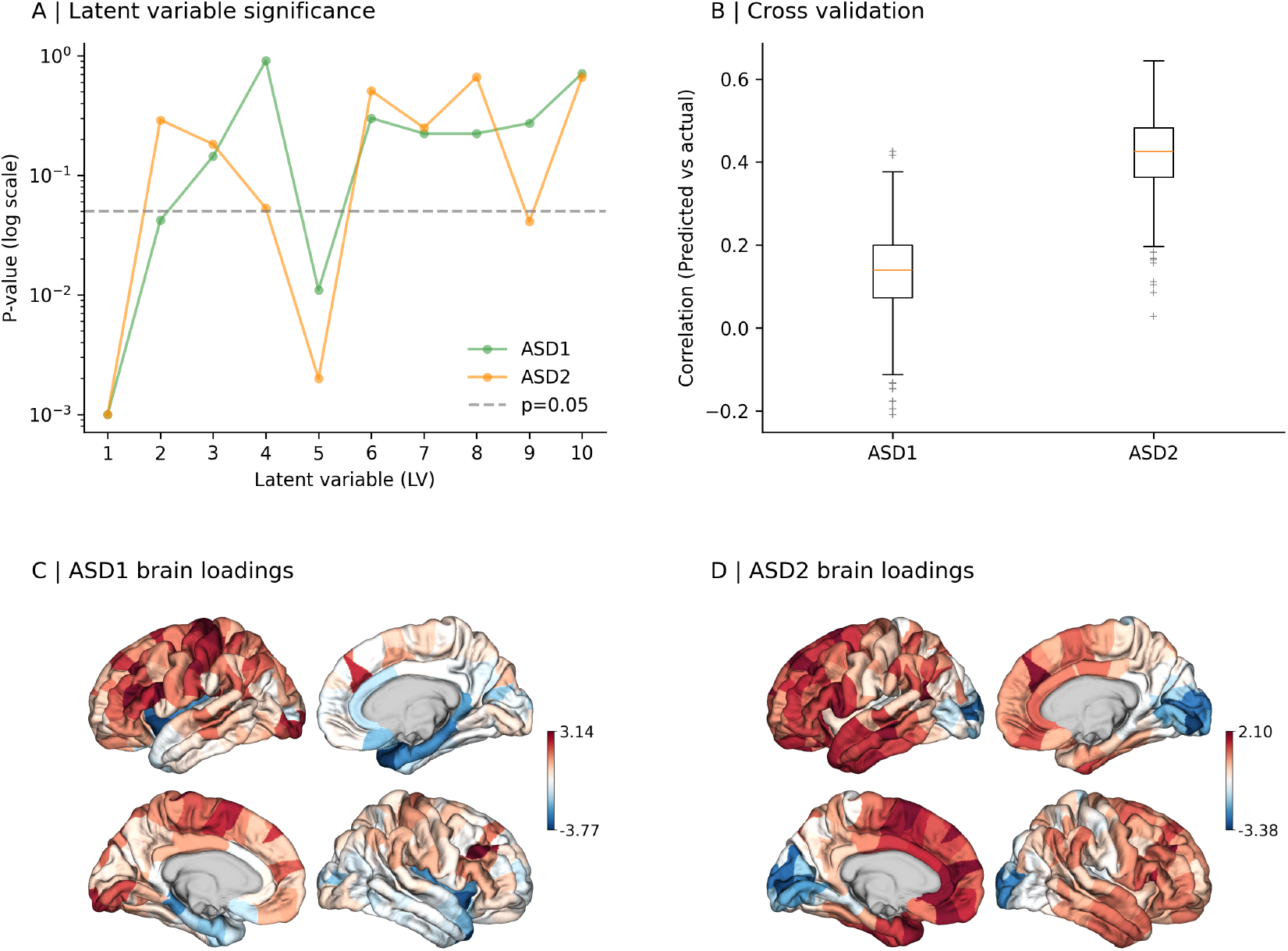
PLSR analysis of gene expression to MIND deviation relationships. (A) P-values for each of the 10 LVs from PLS regression analysis are shown on a logarithmic scale. The dashed gray line indicates the p=0.05 significance threshold. (B) Boxplot summarizing distance-dependent cross-validation performance for each group (median = orange line; box = IQR; whiskers = 1.5×IQR). Z-scored PLS brain region loadings for ASD1 (C) and ASD2 (D), displayed on brain surface maps. Red regions indicate positive loadings (regions where gene expression patterns are positively associated with MIND deviations), while blue regions indicate negative loadings (regions where gene expression patterns are negatively associated with MIND deviations).

**Figure S15.**
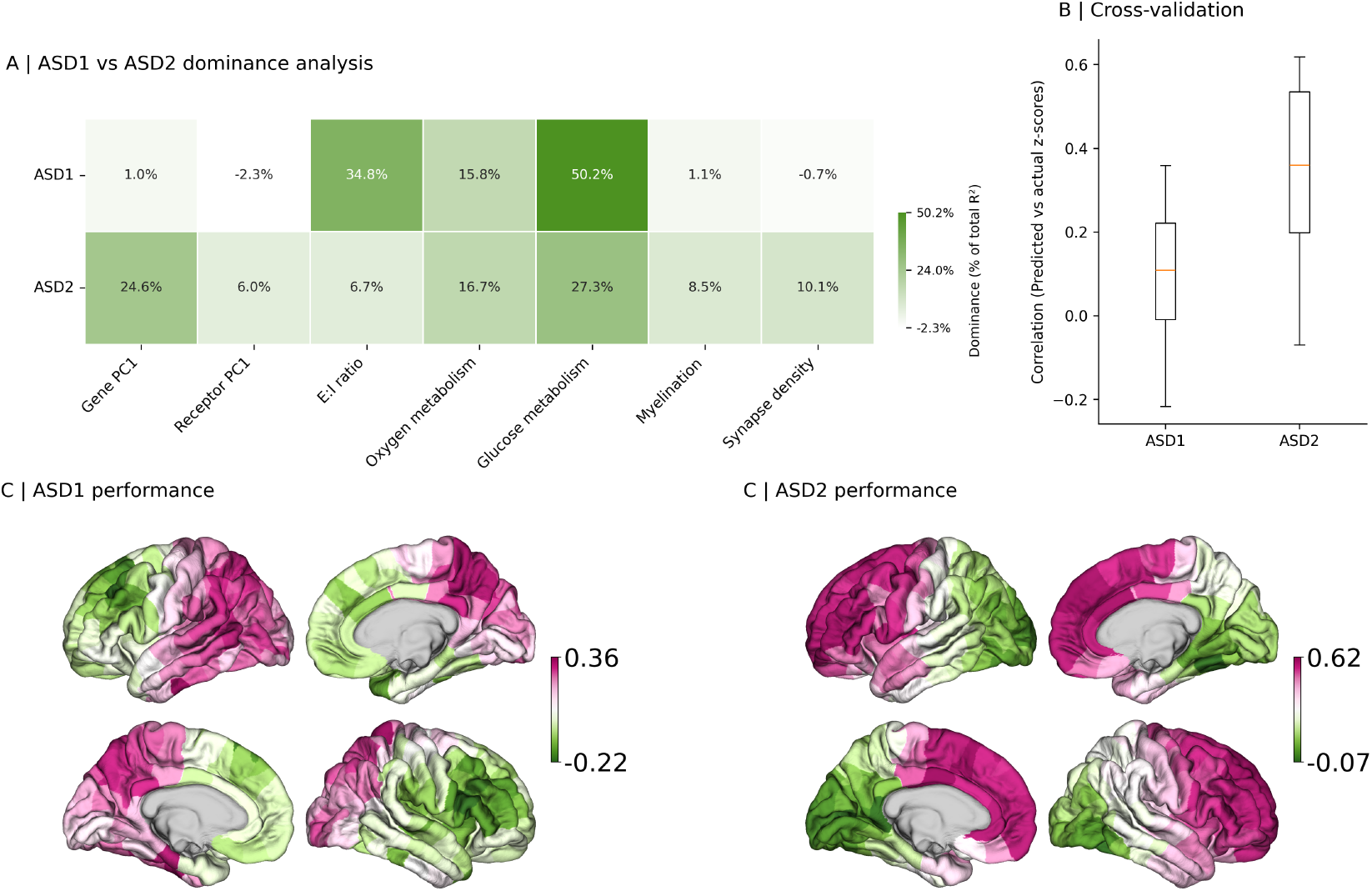
Dominance of biological attributes predicting regional MIND deviations. (A) Heatmap of dominance contributions for seven biological predictors. Each cell reports the absolute dominance (top) and the same value expressed as percent of total adjusted R2 (parentheses). Color encodes percent of total R2. (B) Boxplot summarizing distance-dependent cross-validation performance for each group (median = orange line; box = IQR; whiskers = 1.5×IQR). (C–E) Cortical surface maps of test-set Pearson r (predicted vs observed) for ASD1 (C), ASD2 (D) and All ASD (E) using the distance-dependent 75/25 split.

**Figure S16.**
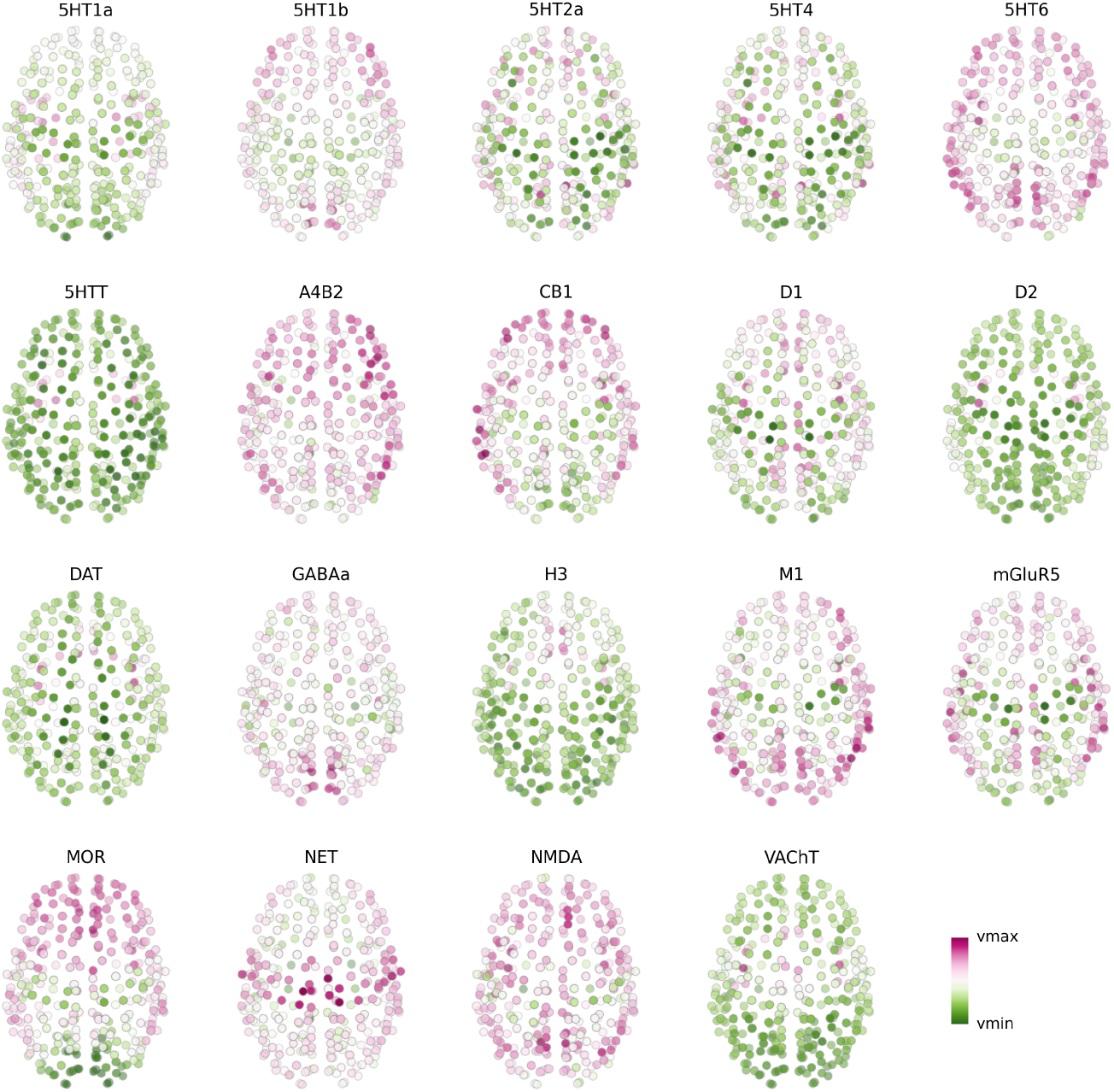
Regional receptor/transporter density maps. Dot-brains show for each of the 19 PET tracers aggregated to the 308 brain regions (color scale indicates relative density).

**Figure S17.**
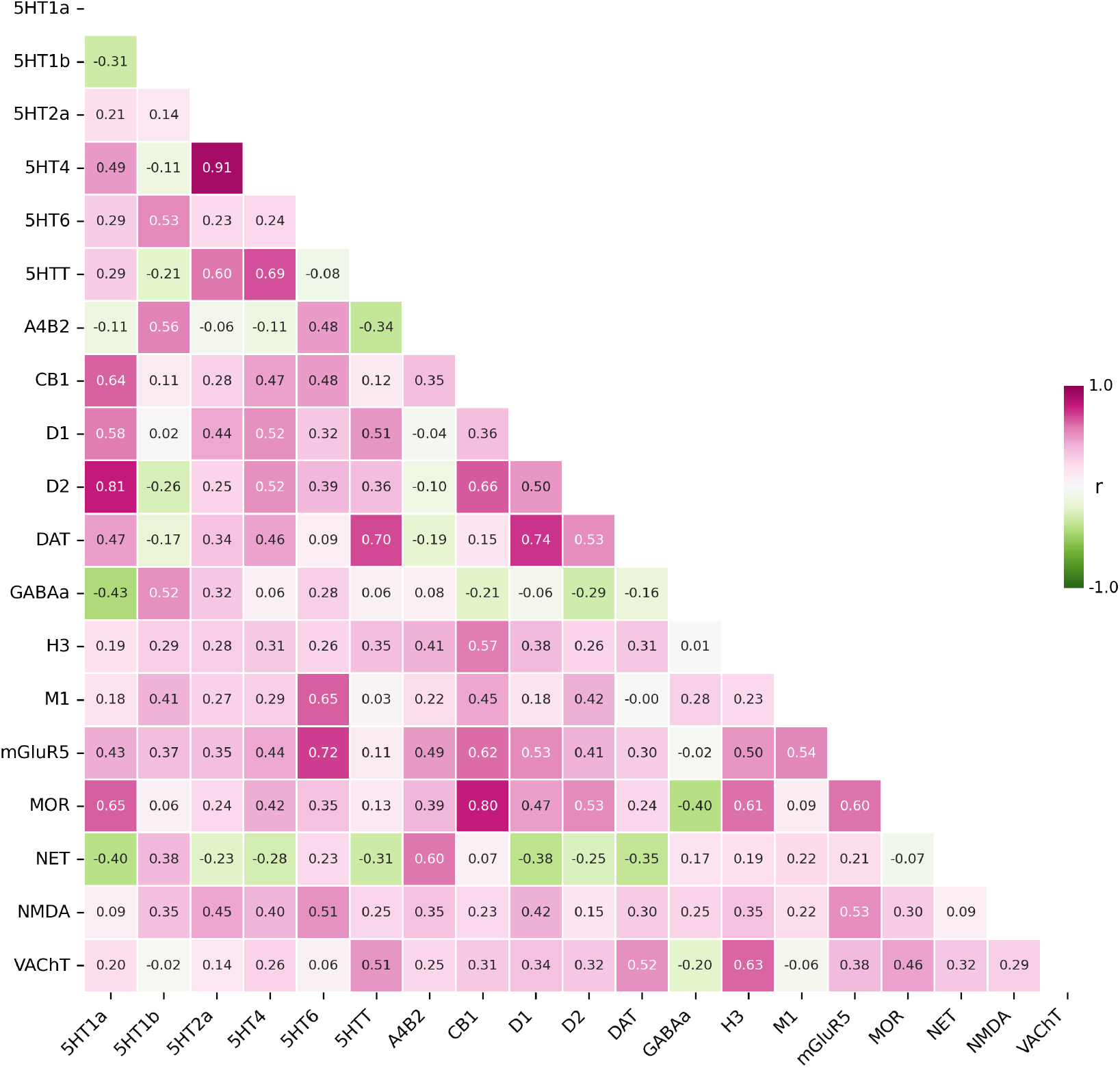
Pairwise correlations among PET-derived receptor/transporter maps. Correlation matrix (Pearson r) across 19 PET-derived molecular targets.

**Figure S18.**
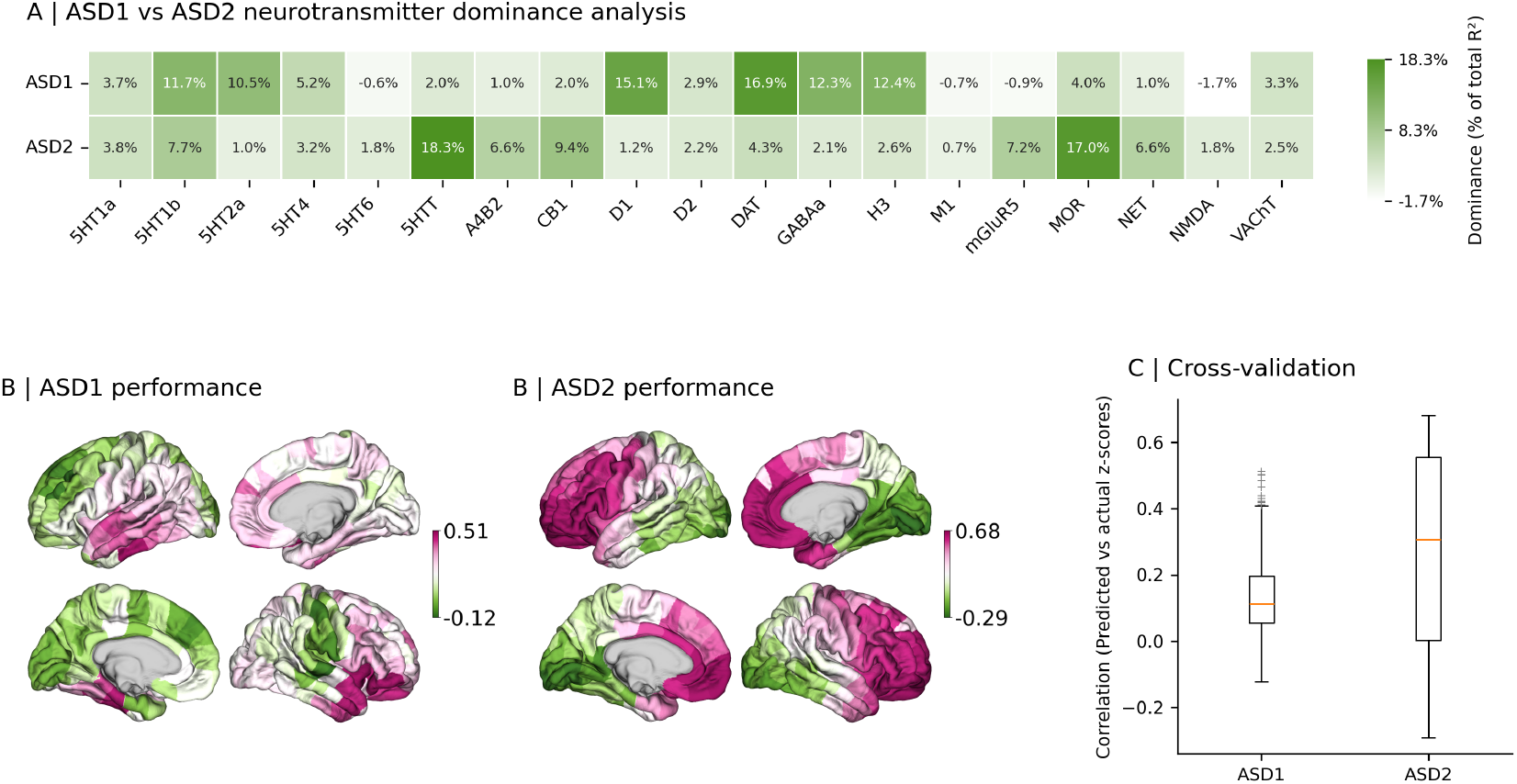
Dominance decomposition for receptor and transporter predictors. (A) Heatmap of dominance contributions for 19 neurotransmitter receptors/transporters. Each cell reports absolute dominance (top) and percent of total adjusted R2 (parentheses); color encodes percent of total R2. (B–D) Distance-dependent cross-validated cortical maps of test-set Pearson r for ASD1 (B), ASD2 (C) and All ASD (D). (E) Boxplot of region-wise test correlations (predicted vs observed) summarizing spatial generalizability.

